# A modular model integrating metabolism, growth, and cell cycle predicts that fermentation is required to modulate cell size in yeast populations

**DOI:** 10.1101/2023.11.25.568635

**Authors:** Marco Vanoni, Pasquale Palumbo, Federico Papa, Stefano Busti, Laura Gotti, Meike Wortel, Bas Teusink, Ivan Orlandi, Alex Pessina, Cristina Airoldi, Luca Brambilla, Marina Vai, Lilia Alberghina

## Abstract

For unicellular organisms, the reproduction rate and growth are crucial determinants of fitness and, therefore, essential functional manifestations of the organism genotype. Using the budding yeast *Saccharomyces cerevisiae* as a model organism, we integrated metabolism, which provides energy and building blocks for growth, with cell mass growth and cell cycle progression into a low-granularity, multiscale (from cell to population) computational model. This model predicted that cells with constitutive respiration do not modulate cell size according to the growth conditions. We experimentally validated the model predictions using mutants with defects in the upper part of glycolysis or glucose transport. Plugging in molecular details of cellular subsystems allowed us to refine predictions from the cellular to the molecular level. Our hybrid multiscale modeling approach provides a framework for structuring molecular knowledge and predicting cell phenotypes under various genetic and environmental conditions.

## INTRODUCTION

The coordination of metabolism, cell growth, and the cell cycle is a central property of all proliferating cells. The budding yeast *Saccharomyces cerevisiae*, whose genetic, molecular, metabolic, and cell physiological understanding is unsurpassed by those of other organisms ^1,2^, is a convenient model system to study this coordination, since many yeast genes involved in essential cellular functions are conserved up to humans, making yeast an attractive experimental model even for human cancer and drug discovery ^3^.

Specific processes within the cell division cycle impose metabolic requirements for energy and biosynthesis of macromolecular precursors, and the metabolic state of individual cells within a growing culture is affected by the cell cycle position. For instance, cells deprived of essential carbon or nitrogen sources arrest at the G0/G1 phase of the cell cycle ^4,5^, and the mobilization of energy stores, such as lipids, trehalose, and glycogen, occurs at the G1/S transition and is controlled by oscillations in Cdk activity ^6-8^. Previous studies have identified interlocked metabolic and cell division cycles ^9,10^, whose phase shifts are a function of environmental conditions ^11-14^. Because metabolic cycles can occur without cell cycle progression, the actions of cell cycle regulatory proteins may only partially explain the cell cycle-specific regulation of metabolism.

The analysis of steady-state cultures can allow us to neglect these intercellular differences. In chemostat experiments, carbon source-specific growth-regulated genes control mitochondrial function, peroxisomes, and synthesis of vitamins and cofactors ^15^. Under these conditions, cells adopt larger sizes and grow faster at higher dilution rates, where metabolism is primarily fermentative ^16-20^. The preference for fermentation over respiration in fast-growing cells is also widespread in higher eukaryotes, including cancer cells ^21,22^. Metabolic fluxes depend on elaborate regulatory processes ranging from signaling events that may modulate the epigenome to control transcription, post-translational modifications, and allosteric regulation of metabolic enzymes. These and other similar events affect metabolism, which sets specific growth dynamics and cell cycle progression. Import of nutrients and their metabolic conversion into building blocks and energy (ATP) sustains yeast cell growth, mainly by net RNA and protein syntheses ^5,23,24^. In turn, cell growth drives the cell cycle ^25,26^. These functions account for the majority of the protein investment in yeast under different nutritional conditions ^27^. Both cell growth and cell cycle progression are controlled by many signaling pathways ^5,25^, including glucose signaling pathways ^28^ and the protein kinase A (PKA)/Tor pathways ^29,30^. Several critical metabolites known to regulate intracellular signaling pathways, such as acetyl-CoA, NADH/NAD^+^, and AMP/cAMP, may trigger growth and cell cycle dynamics. For instance, 90% of the glucose-induced changes can be recapitulated by the activation of PKA or the induction of protein kinase B (PKB; Sch9) ^31^.

The physiological state of a yeast population depends on subtle and sophisticated regulation between two relevant bioprocesses: growth and the cell cycle ^32^. During balanced exponential growth, the physiological parameters of a population remain constant ^33^: these include temporal parameters (such as the mass duplication time, the time required to duplicate DNA, and the fraction of cells in different phases of the cell cycle) and biochemical parameters, such as the average cell size or the distribution of cell size (or protein content) within the growing population. Yeast cells reproduce asymmetrically by budding, yielding a larger parent and smaller daughter at each division (Fig. 1A). To maintain size homeostasis, daughters spend more time in the G1 phase than their cognate parent cells before reaching the critical size required to enter a new cell cycle ^26,34^. Thus, each round of division produces increasingly larger parents and daughters with shorter division times, giving rise to wide heterogeneity in the structure of yeast cell populations ^35,36^. Because the size distribution of asynchronously growing yeast cells is related to protein (biomass) accumulation during the cell cycle, the law of cell division of the population, and its structure, the protein (size) distribution acts as a fingerprint for each condition of balanced exponential growth (Fig. 1B). ^18^

**Figure 1.**
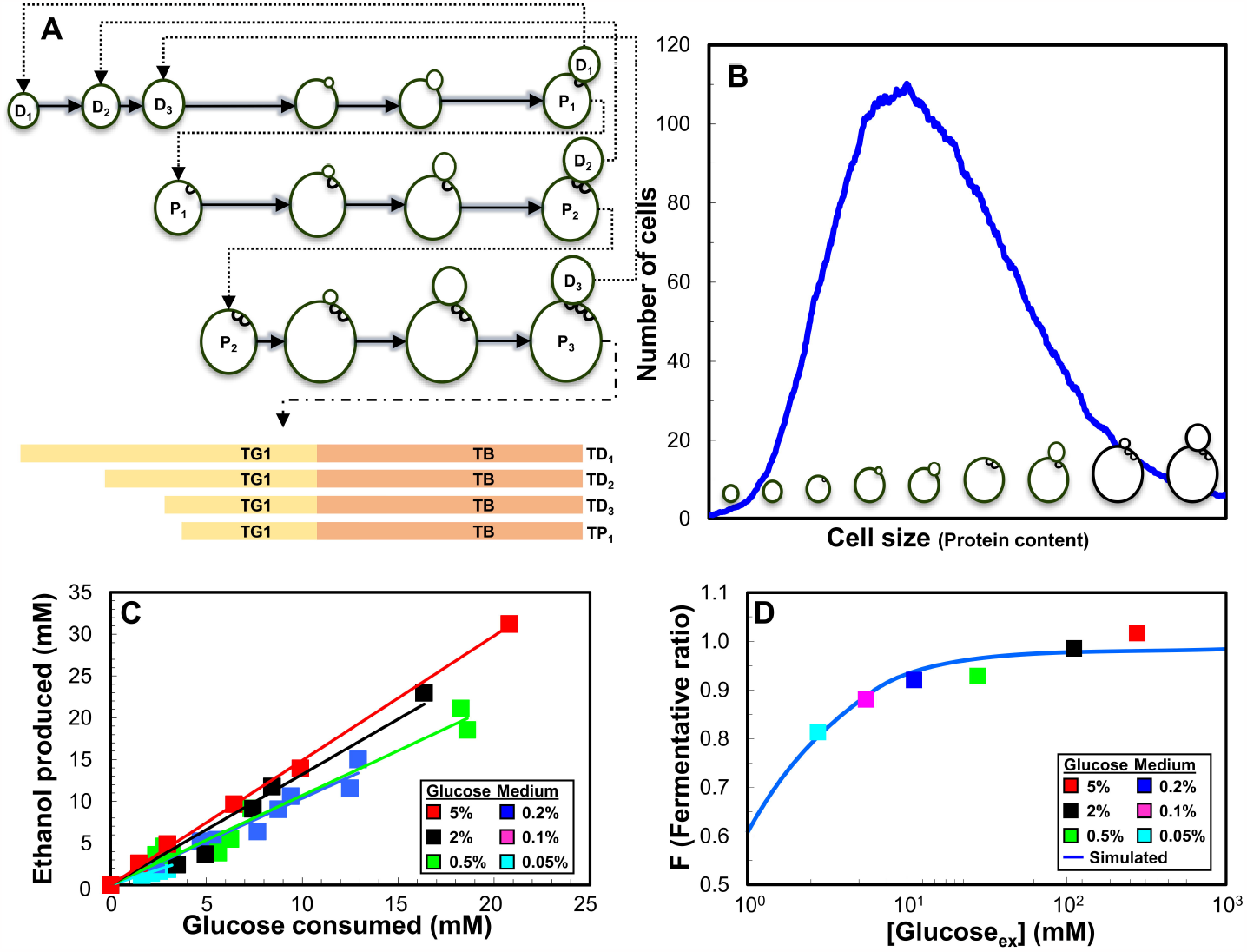
Nutritional modulation of the yeast cell cycle. (A)Schematic view of the cell cycle with cycle times for parents (P) and daughters (D) with different genealogical ages. (B)Cell size (protein content) distribution. (C)Glucose consumption and ethanol production in the wild-type strain in exponential growth. (D)The fermentative ratio F plotted as a function of the external glucose concentration in the growth medium. Colored dots represent experimental data points measured at different glucose concentrations (ranging from 0.05% to 5% glucose); the continuous blue line represents the best fit of the experimental data based on the saturating function reported by Equation (34) of the STAR Methods.

A bottom-up approach has been used to construct a genome-wide whole-cell model for the small parasitic bacterium *Mycoplasma genitalium* ^37^, and a similar approach has been used to model *Escherichia coli* ^38,39^. In this study, we applied a top-down systems biology approach to understand how metabolism controls the growth and division of eukaryotic cells under a given set of nutritional conditions ^40,41^. Starting from a mathematical model that links metabolism, growth (i.e., ribosome and protein accumulation), and cell cycle progression in an average yeast cell ^42^, we present a population model that quantitatively describes the temporal and cellular parameters of yeast populations, such as the protein distributions and average RNA and protein content. Careful tuning of a few selected input parameters allows the recapitulation of alterations in protein distributions and temporal parameters in different environments or genetic backgrounds, enabling the identification of cellular function(s) affected by genetic or environmental perturbations. We validated the model prediction that glucose metabolism (specifically, the ratio between glucose fermentation and respiration) drives nutritional modulation of cell size in exponentially growing yeast populations. Finally, we proved that the model can act as a scaffold for molecularly detailed sub-models, allowing its use to refine predictions from the cellular to the molecular level.

## Results

### A coarse-grained model linking metabolism to cell growth and the cell cycle describes the structure of exponentially growing yeast populations

We recently presented the integrated Metabolism Growth and Cycle (*iMeGroCy*) model that quantitatively describes the complex interrelations connecting metabolism, growth, and the cell cycle ^42^. In this study, we extended the original Metabolism & Growth (*MeGro*) module of *iMeGroCy* by allowing biomass production from ethanol respiration (Fig. 2A) and extending the model to cell populations. The model is described in detail in the *STAR Methods*.

**Figure 2.**
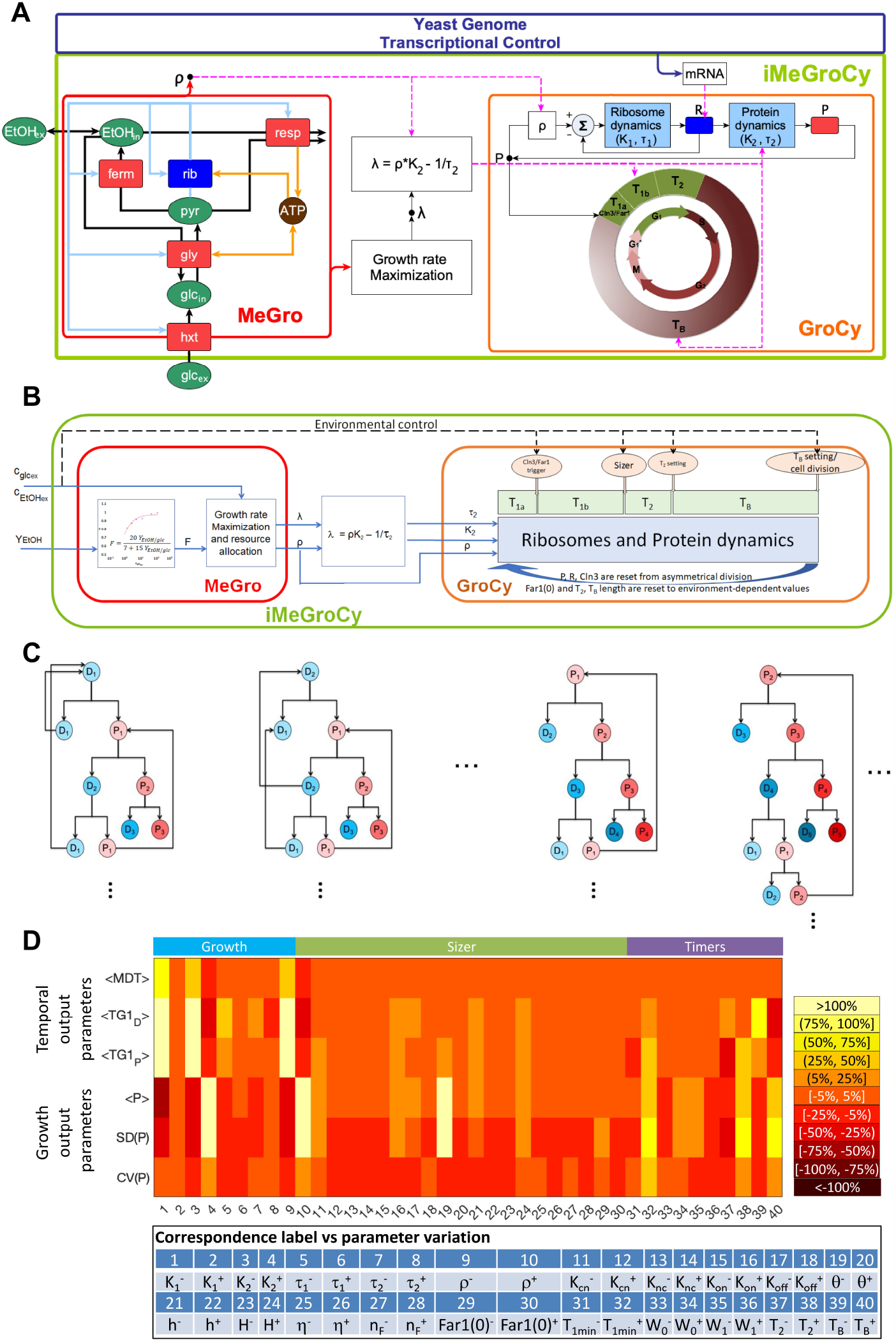
Structure and utilization of the integrated Metabolism Growth and Cycle model. (A)The *iMeGroCy* model (green block) includes two main modules: *MeGro* (red block) and *GroCy* (orange block), which are interconnected by dashed magenta arrows. Black solid arrows: metabolic conversions; cyan solid arrows: protein synthesis; orange solid arrows: ATP production/consumption; blue rectangles: ribosomes (rib); red rectangles: other protein classes - glucose transporter (hxt), glycolytic enzymes (gly), enzymes of the fermentative pathway (ferm), enzymes of the respiratory pathway (resp); green ellipses: metabolites - external glucose (glc_ex_), internal glucose (glc_in_), pyruvate (pyr), internal and external ethanol (EtOH_in_ and EtOH_ex_ respectively); brown circle: energy carrier - ATP. (B)*iMeGroCy* block diagram highlighting the wiring among the component modules. (C)Pedigree map populations derived from germinal lines starting from different progenitor cells. (D)Graphical summary of alterations in major output growth and temporal parameters by changing input parameters of the *GroCy* module.

The updated version of *MeGro* includes lumped glycolytic and gluconeogenic pathways, a fermentation pathway, and a respiratory pathway, which can be fed by either pyruvate or ethanol, a glucose transporter unit, a membrane transport mechanism for ethanol, and ribosomes devoted to biomass production. In addition, ethanol derived from the outside can be respired or used in a simplified gluconeogenic pathway, allowing *MeGro* to sustain the synthesis of building blocks without extracellular glucose.

The Growth & Cycle module (*GroCy*) links cell growth to the cell cycle. Cell growth is modeled by an Ordinary Differential Equations (ODE) system describing ribosome and protein dynamics (synthesis and degradation), and the interaction between Cln3 (the most upstream G1 Cdk1 activator that increases proportionally to cell mass) and Far1 (an inhibitor of the Cln3/Cdk1 complex). The cell cycle is modeled as a series of consecutive timers. Timers T_1A_, T_1B_, and T_2_ encompass the G_1_, unbudded phase. T_B_ corresponds to the budded phase, which includes the S, G_2_, M, and G_1_* phases, of which the latter corresponds to the period that separates nuclear and cell division.

Fig. 2B shows the wiring between the two modules of *iMeGroCy. MeGro* acts as a parameter generator, receiving two inputs for each growth condition: the external carbon source concentration ( *C*_*glc*_ or *C*_*ETOH*_) and the fermentative ratio F, calculated from the experimental ethanol/glucose yield Y_EtOH_ for glucose-grown cells (see *STAR Methods*) or set to 0 for ethanol-grown cells. Using these inputs, *MeGro* maximizes the exponential growth rate λ and allows calculation of the ribosome and protein dynamics parameters fed to the *GroCy* module (see *STAR Methods* for details). In addition, growth and cell cycle parameters not directly provided by *MeGro* are directly input by the user (see below and the *STAR Methods* for further information on parameter tuning).

Ribosome and protein dynamics control the Cln3/Far1 ratio. Nuclear Cln3 binds to its inhibitor Far1, forming the Cln3/Far1 complex at birth. As newborn cells grow, total Cln3 increases proportionally with the overall cell protein content, thus enabling nuclear Cln3 to overcome Far1, which is more efficiently degraded when unbound from Cln3. The end of the T_1A_ timer (which starts at cell birth) is reached when Cln3 is more abundant than the Cln3/Far1 complex, triggering T_1B_, whose length is inversely proportional to the protein content of the cell. The T_B_ timer begins after period T_2_ of fixed duration. The cell simulation ends when the T_B_ timer elapses. Ribosomes and proteins are asymmetrically divided between the newborn daughter, which receives the mass synthesized from the onset of budding, and the parent cell, which receives the mass of the original parent at the start of budding; the same asymmetrical division occurs for the cytosolic molecular players. In contrast, nuclear Far1, nuclear Cln3, and any remaining nuclear Cln3/Far1 complex are symmetrically divided at the onset of the G_1_* phase (the last 20% of the T_B_ timer), and Far1 was reset to its starting value (see ^42^ and the *STAR Methods* for details).

We used a pedigree structure to describe the exponential growth of the yeast populations, following the destiny and recording parameters of cell clusters originating from a single founder cell (Fig. 2C). After each cell division, the model parameters were randomized, as described in the *STAR Methods*, to simulate cell-to-cell biological variability. After sufficient generations, the simulated populations reached a steady state, in which the representative parameters of the population, including the average protein content and fraction of budded cells, remained stable over time (Fig. S1A-D). The final population used in this study was obtained by mixing 10 clusters, each containing at least 104 cells. Fig. S1E shows an example of the protein distribution obtained through this procedure.

To assess the effect of each input parameter on the output growth and temporal parameters, we changed a single input parameter while keeping the remaining input parameters constant. The monitored temporal parameters constitute the average value, standard deviation, and coefficient of variation of the protein distribution (<P>, SD(P) and CV(P), respectively) while the temporal parameters constitute the mass duplication time of the population (MDT) and the length of the G_1_ phase of parent and daughter cells (TG1_D_ and TG1_P_, respectively). Briefly, each input parameter has been moved “forward” and “backward” in comparison with its nominal value, with the number and amplitude of the variations chosen on the basis of an *a priori* evaluation of the biological meaning of each parameter and the limits assigned by the onset of computational problems. For each parameter setting (resulting from each parameter variation), exponential growth of the yeast population was simulated (up to ∼50000 cells), linking the *GroCy* input parameters to the quantitative features of the cell population. As a representative example, Fig. S2 shows a sensitivity analysis of parameter T_2_. The maximal alteration in the temporal output and growth parameters caused by altering each input parameter is plotted in Fig. 2D. A value of 100% indicates that the output parameter is doubled in comparison with the reference value. The resulting sensitivity map (Fig. 2D) provides a reference database linking the input model parameters to the quantitative output features of the virtual cell populations. Perturbation of a large set of parameters had a limited effect on the tested output parameters, indicating that the model was robust against parameter alterations.

### Metabolism drives the coordination between cell growth and division

Different nutrients and growth conditions affect the cell size and cell cycle parameters, including the growth rate, budding index, and length of the budding phase (Table S1A) ^17,43-45^. Yeast cells can use ethanol only through respiration, whereas respiration or fermentation, which produce ethanol as a byproduct, can metabolize glucose. To investigate the link between metabolism and cell size modulation, we measured ethanol production as a function of glucose consumption (Fig. 1C). The lower the glucose concentration in the growth medium, the lower the ethanol produced for each unit of consumed glucose (Fig. 1D), indicating increased respiratory utilization of sugar.

As outlined above, modification of only a few parameters may produce yeast populations with significantly different properties. Successful prediction of the output parameters of cell cultures grown in different growth media would indicate that the model can correctly identify the crucial parameters that are affected by the experimental conditions (or genetic mutation; see the next subsections). To adapt the model input parameters to different glucose concentrations, we used the generated knowledge base (Fig. 2D) and manually adjusted a few parameters following the procedure outlined in the block diagram in Fig. 3A. *MeGro* (see Fig. 2B) uses the experimental fermentative ratio F to compute the parameters of protein dynamics (K_2_, ρ) for cells grown in glucose as inputs for the *GroCy* module. By definition, F is zero for ethanol-grown cells. A few input parameters of the cycle module (*i*.*e*., the initial level of Far1 and the average lengths of T_1_ and T_B_) were manually tuned according to the growth conditions ^42,46,47^, as highlighted by the dashed black line in Fig. 2B. For example, the experimental data obtained for poorer media revealed a substantial reduction in the mean value and standard deviation of the protein content, as well as a considerable increase in the mass duplication time and the G_1_-phase length, in comparison with the reference condition, i.e., glucose 2% (Table S1A). Fig. 2D indicates that these population features are achievable by reducing K_2_, ρ (that simultaneously produces a reduction in <P> and SD(P) and an increase of <MDT>, <TG1_P_>, <TG1_D_>) and W_0_ (that accounts for a reduction in <P> and SD(P) and does not affect <MDT>, <TG1_P_>, and <TG1_D_>). The input and output parameters used in the simulations are listed in Tables S2 (*MeGro* input parameters), S3 (*iMeGroCy* input parameters) and S4 (*iMeGroCy* population output parameters).

**Figure 3.**
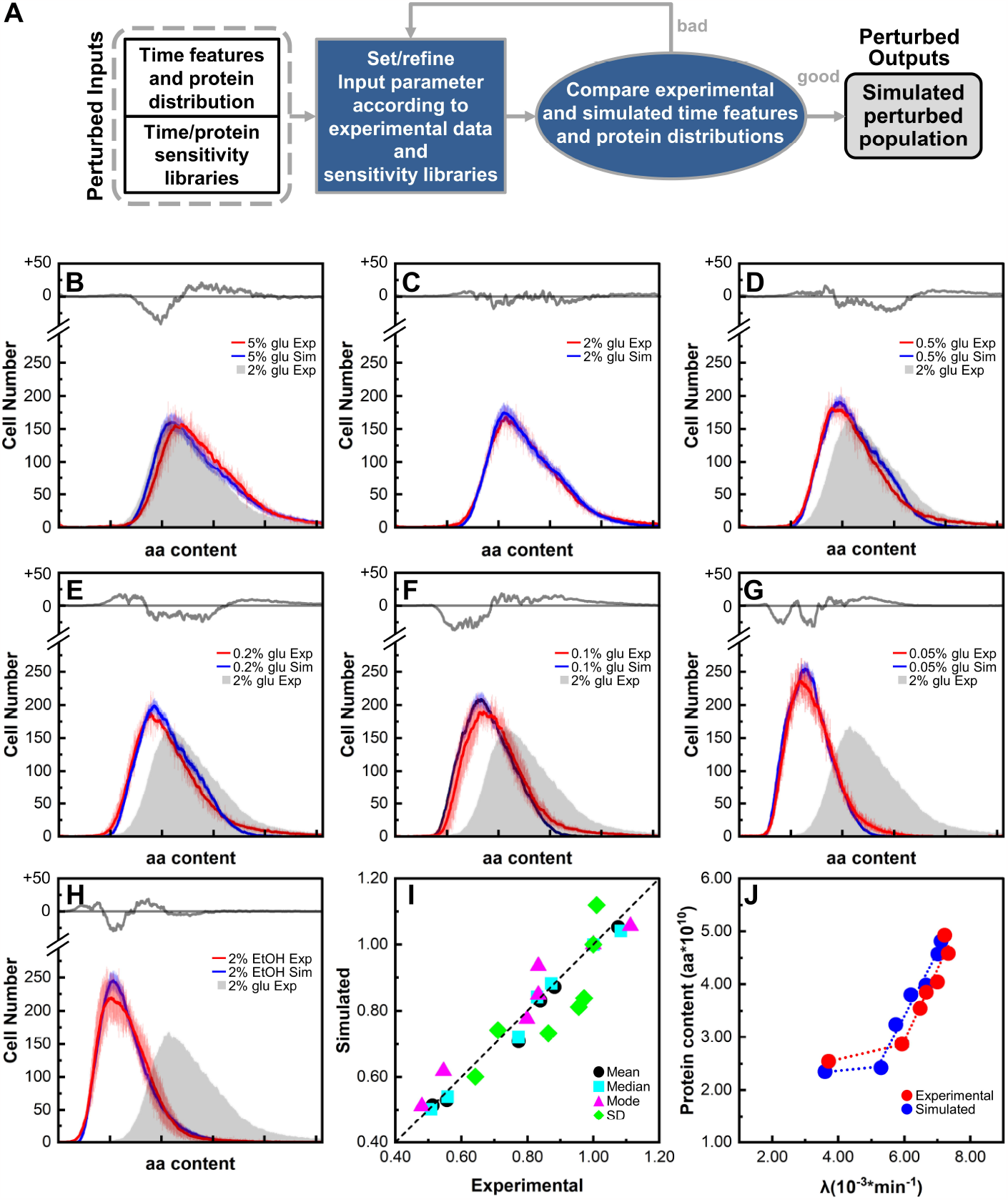
A coarse-grained model describes nutritional modulation of growing yeast populations. (A)Flow chart of the identification procedure for mutational and environmental perturbations. (B-H) Comparison of the experimental (red lines) and simulated (blue lines) protein distributions for wild-type cells grown in media supplemented with different glucose concentrations or ethanol. Mean (blue, red) ± SD values (light blue and light red) are shown. In each panel, the gray line in the upper insert represents the difference between the experimental and simulated values. The experimental distribution of cells grown in 2% glucose (filled gray) is also reported as a reference. (I)Correlation between protein distribution parameters of simulated and experimental yeast populations. (J) Growth rate (λ, min^-1^) vs. average protein contents (P, aa) in media supplemented with different glucose concentrations or ethanol. Experimental (red) and simulated (blue) values (mean ± SD) are reported.

As a preliminary validation step, we checked whether our model could simulate different nutritional conditions using roughly the same set of parameters because the MeGro optimization module reacts differently to different environments. Fig. 3B-H shows the experimental (red) and simulated (blue) protein distributions for populations of the wild-type CEN.PK strain proliferating in media supplemented with either ethanol or different amounts of glucose (from 0.05% to 5%) as the carbon source. The reported data include the average ± standard deviations of the experimental (red) and computational (blue) cell numbers with a given protein content. Despite minor differences between the observed and simulated distributions (drawn in the upper part of panels B-H using the same Y-scale as the protein distributions), the simulated distributions closely matched the experimental distributions. Most differences were confined within one standard deviation, strengthening the ability of the model to faithfully reproduce distributions of cell protein content (used here as a proxy of cell size) ^4,48^ of yeast populations in exponential growth. The gray distribution (the experimental protein distribution of cells grown in the reference condition, i.e., 2% glucose) provided immediate visual feedback on the ability of the simulated distribution to capture the nutritional modulation of cell size (protein content). Accordingly, the mean, median, mode, and standard deviation of the corresponding experimental and simulated distributions showed excellent correlations (Fig. 3I). Finally, Fig. 3J reports the average experimental (red) and simulated (blue) protein content as a function of the growth rate (λ, min^-1^). For both experimental and computational distributions, glucose concentration strongly modulates the average protein content, which shows a steep linear correlation with the growth rate λ (Fig. 3J). Conversely, growth in ethanol-supplemented media significantly decreases λ, which nearly halves from 0.05% glucose to ethanol (Fig. 3J; Table S1A), but results in only a further marginal reduction in P (ca. 10%). In conclusion, our results indicate that by reducing the glucose concentration in the growth medium, metabolism progressively shifts toward respiration, and that the *iMeGroCy* model captures not only the change in growth rate but also the overall size distribution in exponentially growing populations. As outlined in the Introduction, the protein (size) distribution is a fingerprint of each condition of balanced exponential growth, and the ability to capture this modulation by tweaking only a few parameters indicates that the simplicity of the model includes the most relevant biological interconnection among metabolism, growth, and the cell cycle.

### Metabolism-driven modulation of cell size

Our data confirmed that glucose can modulate cell size in a dose-dependent manner as a function of the growth rate: increasingly higher sugar concentrations in the growth medium support faster growth rates and larger sizes (Fig. 3; Table S1A) ^17,24,44,45^. However, the existence of a linear correlation between the yield of ethanol and the protein content of the population (P, Ps, and P0) suggests that the type of energy metabolism (respiratory, respiro-fermentative, and fermentative) of the cell modulates both growth rate and size (Fig. S3A).

In glucose-limited continuous cultures, the cell size has been consistently shown to be small and relatively constant at low growth rates when only respiration is active. In contrast, it increases steeply at faster growth rates when cells switch to fermentation, resulting in ethanol production ^17,43^. Furthermore, cell size increases even at low growth rates if the cells are forced to shift from respiratory to fermentative metabolism ^17^.

Accordingly, our model predicted that for any given fixed value of the fermentative-to-respiratory ratio (F), ranging from 0 (fully respiratory metabolism) to 1 (pure fermentation), the dose-dependent modulation of cell size (estimated by the parameter P, the average protein level of the population) by external glucose was substantially lost for F values ≤ 0.8, with the protein content remaining evenly small regardless of the external glucose concentration (Fig. 4A). The nutritionally dependent modulation of the protein content occurs only when glucose metabolism is strongly shifted toward fermentation, being abolished when cells are “forced” to adopt a respiratory metabolism.

**Figure 4.**
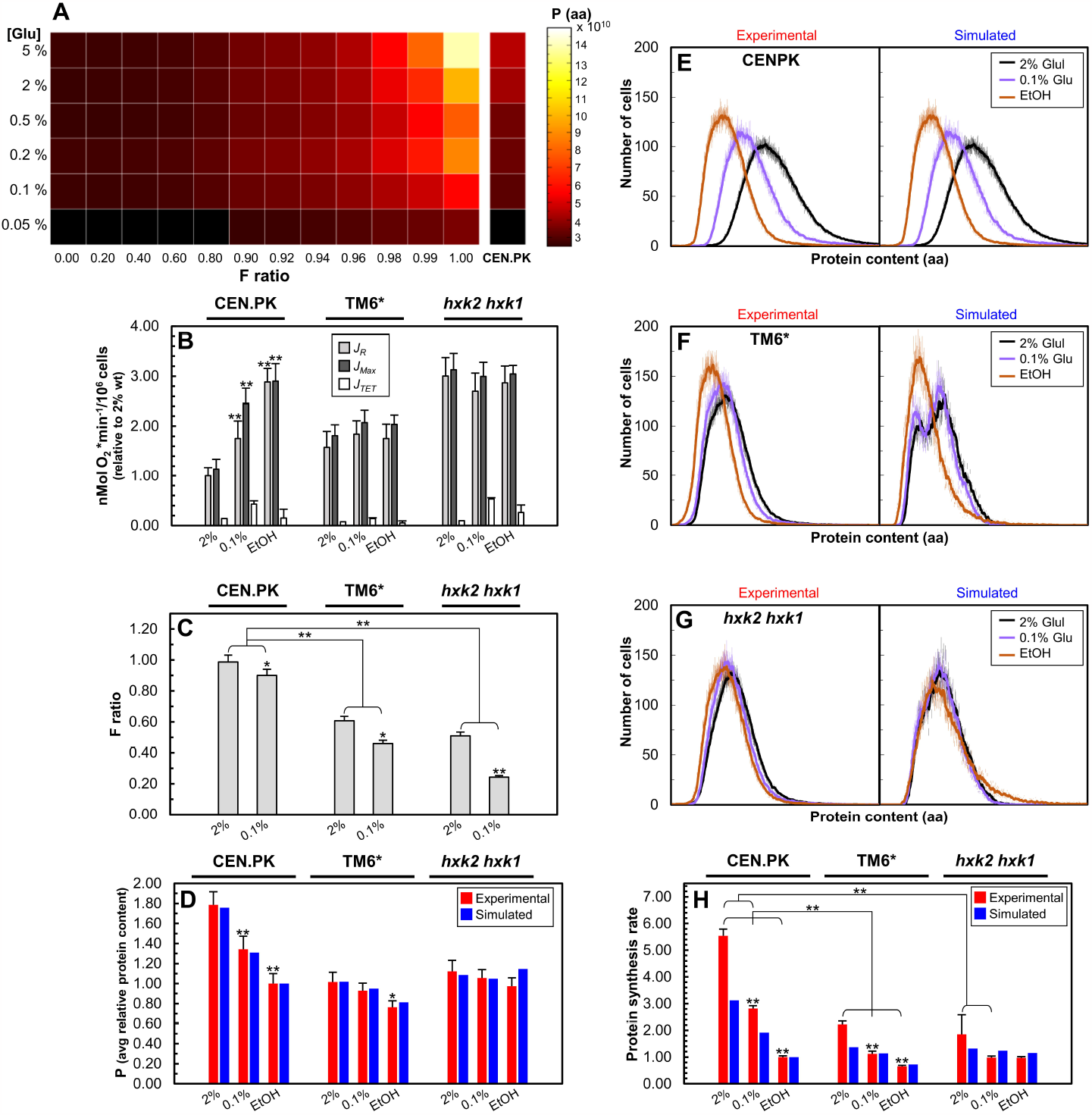
A coarse-grained model describes metabolism-driven modulation of cell size. (A)Mean protein content of simulated *iMeGroCy* yeast populations obtained for different external glucose environments (ranging from 0.05% to 5%) and fermentative conditions (F ratio in [0, 1]). The *iMeGroCy* parameters were set according to the values for the wild-type CEN.PK strain reported in Table S3. The CEN.PK column reports the mean protein content of the *iMeGroCy* population in the same glucose environment, obtained by setting F equal to the experimental value measured for CEN.PK. (B)Respiratory parameters determined for the wild-type CEN.PK strain and TM6* and *hxk2 hxk1* respiratory-deficient mutants. Basal respiration rate (*J*_*R*_), uncoupled respiration rate (*J*_*MAX*_), and non-phosporilating respiration (*J*_*TET*_) during growth in 2% glucose, 0.1% glucose, and ethanol media are shown. Data are relative to wild-type oxygen consumption in 2% glucose medium. Values are mean ± SD from three independent biological replicates. Statistical significance: *p < 0.05, **p < 0.01, Student’s t-test. (C)The F ratio was experimentally determined for wild-type and respiratory-deficient mutants under different growth conditions. Values are mean ± SD from three independent biological replicates. Statistical significance: *p < 0.05, **p < 0.01, Student’s t-test. (D)Experimental (red bars) and simulated (blue bars) average protein contents in the wild-type and respiratory-deficient mutants under different growth conditions. Values are mean ± SD from three independent biological replicates. Statistical significance: *p < 0.05, **p < 0.01, Student’s t-test. (E-G) Experimental and simulated intracellular protein content distributions in the wild-type and respiratory-deficient strains obtained under different growth conditions. Mean ± standard deviation (SD) values are reported. (H) Experimental (red bars) and simulated (blue bars) protein synthesis rates for wild-type and respiratory-deficient mutants obtained under different growth conditions. Data are relative to the findings for the wild-type strain cultivated in ethanol medium. Mean + SD values from at least three biological replicates are reported. Statistical significance: *p < 0.05, **p < 0.01, Student’s t-test.

To validate these predictions, we analyzed the behavior of two mutants defective in glucose metabolism: the TM6* strain, which exhibits strongly reduced sugar uptake capacity (Fig. S3I), ^49^ and the *hxk2 hxk1* strain, which lacks two of the phosphorylating isoenzymes that catalyze the first glycolytic step (Fig. S3J) ^50,51^. Cell size parameters (P, Ps, and P0) were monitored during balanced growth in media supplemented with high (2%) or low (0.1%) glucose levels and ethanol and used as a reference for full oxidative metabolism (Table S1B, Fig. S3B-C, F-G). We consider that the nutritional modulation of cell size is fully operative when the ratio of protein content during growth in glucose vs. ethanol is approximately 1.8. In contrast, it is lost when this ratio is close to 1.0. Table S5 shows the parameters altered in the simulations of the TM6* and *hxk1 hxk2* mutants.

In contrast to their wild-type isogenic counterparts, both mutants primarily adopted respiratory metabolism to meet their ATP demands under all tested growth conditions, as suggested by their constitutively high oxygen consumption rates (Fig. 4B) and the concomitant decrease in F ratio values (Fig. 4C) relative to their isogenic wild-type counterparts. This behavior is likely the result of reduced glycolytic flux or relaxation of their glucose-repression mechanisms ^49,50^. Most importantly, the carbon source-dependent modulation of cell size was substantially compromised in both the TM6* and *hxk2 hxk1* mutants (P_2%Glu_/P_EtOH_ = 1.330 and 1.152, respectively). As predicted by our model, under all tested growth conditions, the mutant cells showed a small size regardless of the available carbon source (Fig. 4D), resulting in nearly superimposable protein distribution profiles (Fig. 4E-G). In addition, our model correctly predicted a significant reduction in the protein synthesis rate (relative to the rate in the wild-type strain grown in 2% glucose medium) in these mutants and wild-type cells grown in 0.1% glucose and ethanol medium (Fig. 4H; Fig. S4A). Note that most parameters were inherited from the wild-type population, indicating that the model simulation pinpointed the specific biological alterations underlying the phenotypes of the mutants.

### The integrated model describes the growth properties of both cell growth and cell cycle mutants

As shown in Fig. 5, we altered the input parameters to successfully fit both the temporal parameters and protein distributions of the mutants involved in metabolism, one of the three functions integrated by the *iMeGroCy* model. Here, we analyzed mutants of two more genes: *rsa1*, a mutant defective in ribosome biogenesis ^52^ (Fig. 5A), and two mutants (Fig. 5B-C) of the *WHI5* gene ^53,54^, which encodes an inhibitor of SBF, one of the key transcription factors required for the G_1_/S transition. Phosphorylation at multiple sites dissociates Whi5 from SBF, allowing the transcriptional activation of SBF target genes. Four specific “functional” sites of Whi5 must be phosphorylated to release Whi5 from SBF ^55,56^. Cells lacking the *WHI5* gene (*whi5Δ* strain) or expressing a Whi5 protein whose functional phosphorylation sites have been mutated to phospho-mimetic glutamate residues (*whi54E* strain; ^53^) are expected to enter the S phase. The model allowed for successful fitting of the protein distributions of the *rsa1, whi5Δ*, and *whi54E* mutants, capturing the reduction in the size of exponentially growing populations ^54,56-59^ (Fig. 5A-C). Table S6 shows the parameters altered for the simulations of the *rsa1* and *whi5* mutants. The model predicted a mild reduction in the rate of protein synthesis for *whi5* mutants and a more severe decrease in *rsa1* mutants, which approximated the rate observed in wild-type cells grown in 0.1% glucose. Experimental data confirmed that *rsa1* mutants and, more so, *whi5* mutants showed a reduction in the protein synthesis rate (Fig. 5D; Fig. S4B).

**Figure 5.**
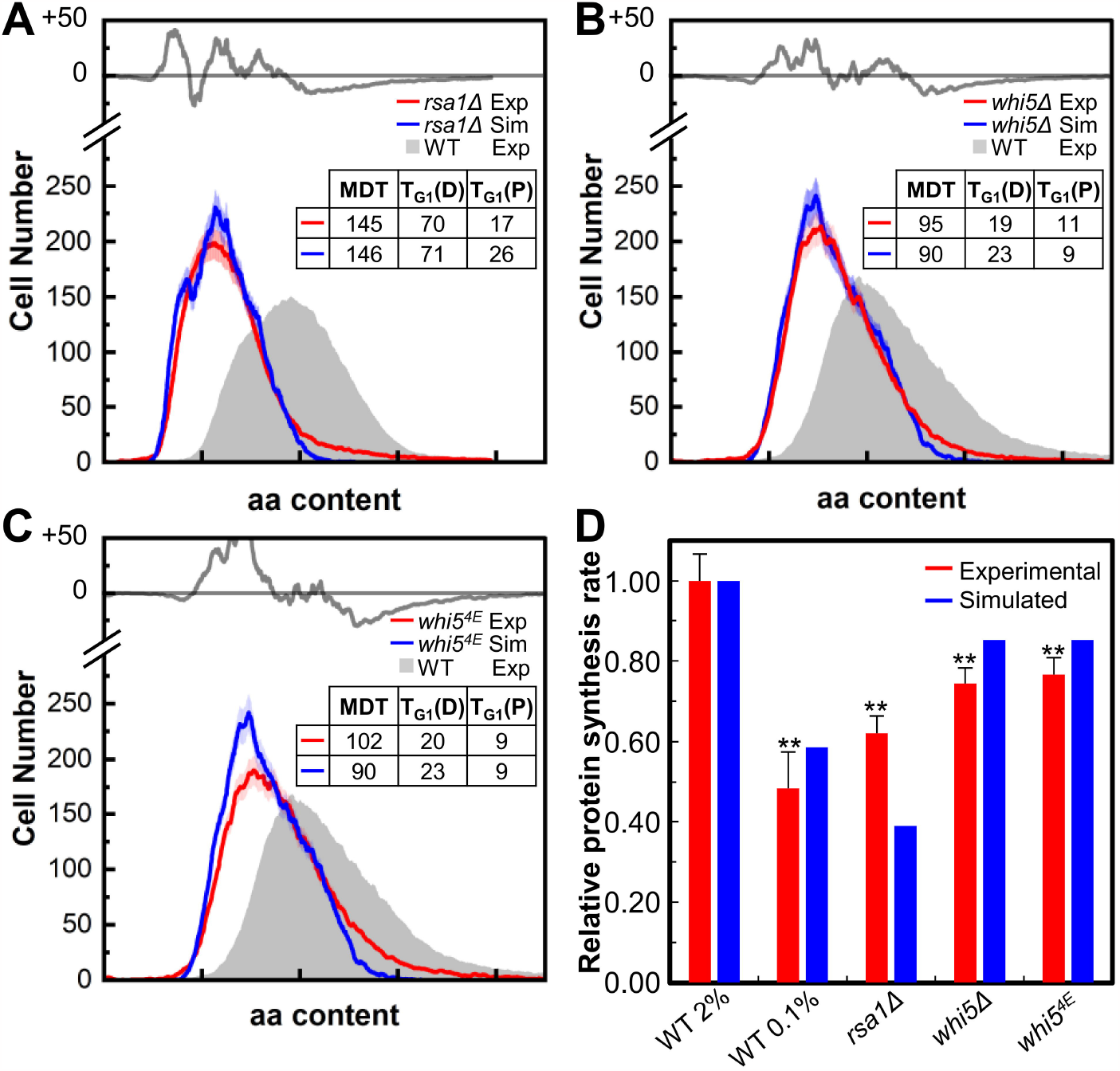
A coarse-grained model predicts the protein distributions of mutants in cell growth and cell cycle. (A-C) Comparison of experimental (red lines) and simulated (blue lines) protein distributions for the mutants *rsa1* (A), *whi5*_Δ_ (B), and *whi5*^*4E*^ (C) grown in glucose 2% media. Mean (blue, red) ± SD values (light blue and light red) are shown. In each panel, the gray line in the upper insert is the difference between experimental and simulated values. The experimental distribution of wild-type cells cultivated in 2% glucose (filled gray) is also reported as a reference. The lower inserts show the temporal parameters for both experimental and simulated populations. (D) Experimental (red bars) and simulated (blue bars) protein synthesis rates for different strains and growth conditions. Data are relative to the wild-type strain cultivated in 2% glucose medium. Mean + SD values from at least three biological replicates are reported. Statistical significance: *p ≤ 0.05 and **p ≤ 0.01, Student’s t-test.

### The integrated model acts as a scaffold for a G_1_/S transition model with molecular resolution

Although the integrated model lacks molecular mechanistic details, it may offer a valid framework for integrating many molecular models of cell subsystems. As a proof of principle, we replaced the Cln3/Far1 trigger and the periods describing the G_1_ phase with a dynamic molecular model of the G_1_/S transition (Fig. 6A ^53^). The resulting hybrid HyG1_S-iMeGroCy model quantitatively described the temporal parameters and protein distributions of the wild-type strain at both high (2%) and low (0.05%) glucose concentrations (Fig. 6B-C; Tables S7 and S8), providing proof-of-principle for the ability of our integrated model to act as a scaffold for molecularly detailed models of selected pathways while retaining the general dynamics of cell functioning. Analysis of the whi5 mutants (Fig. 5) showed excellent agreement between the experimental and simulated temporal and growth parameters. The agreement between the experimental and computational protein distributions was excellent for the *whi5Δ* mutant and good for the *whi54E* strain (Fig. 6D and 6E, respectively), confirming the predictive power of the hybrid model.

**Figure 6.**
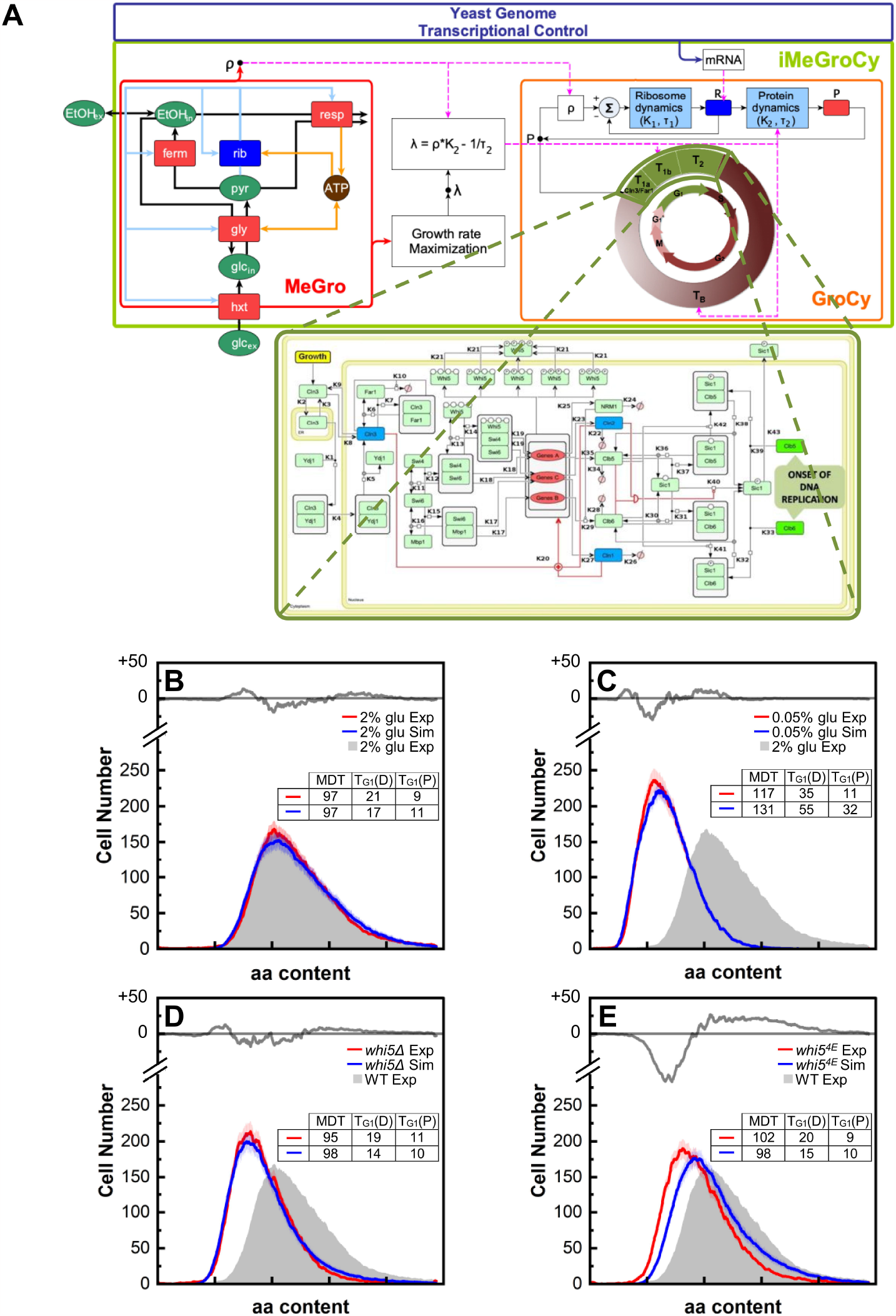
A coarse-grained model acts as scaffold for a molecularly detailed module of the G_1_/S transition. (A) The G_1_/S transition molecular module was plugged into the *iMeGroCy* model to replace Timers T_1_+T_2_. (B–E) Protein distributions predicted by the hybrid *Hy-iMeGroCy* model (blue lines), compared to the related experimental distributions (red lines), for wild-type strains grown in 2% (B) and 0.05% (C) glucose media, and for mutants *whi5*_Δ_ (D) and *whi5*^*4E*^ (E). Mean ± SD values are shown. In each panel, the gray line in the upper insert represents the difference between the experimental and simulated values. The experimental distribution of wild-type cells cultivated in 2% glucose (filled gray) is also reported as a reference. Lower inserts show the temporal parameters for both experimental and simulated populations.

## DISCUSSION

Metabolism, cell growth, and the cell cycle account for the majority of protein investments under different nutritional conditions in yeast ^27^. Metabolism extracts energy from nutrients and sustains yeast cell growth, mainly through net RNA and protein syntheses ^5,23,24^, which control the cell cycle ^25,26^. In this study, we constructed and validated a coarse-grained multilevel dynamic model that combined cellular and population layers to accurately predict quantitative physiological properties (protein distributions and temporal parameters) in response to both genetic variants and the glucose concentration in the growth medium. During steady-state growth, cell size was set almost exclusively by the type of metabolism, with sensing at a higher hierarchical level to fine-tune the metabolism-growth-cell cycle interconnections. Our integrated metabolism, cell growth, and cycle model supports this view because it quantitatively accounts for the temporal and growth parameters of wild-type cells grown at different glucose concentrations or in the presence of ethanol (Fig. 3).

A sensitivity analysis based on the population model predicted that nutritional modulation of cell size requires the ability to ferment glucose. The analysis of two non-fermenting mutants (*hxk2 hxk1* and TM6*, which are defective in glucose phosphorylation and transport, respectively) validated this prediction. The model also described alterations in population structure originating from mutations in growth (*rsa1*, defective in protein synthesis) and cell cycle (deletion and point mutations in the cell cycle inhibitor Whi5). Modifying only a few selected parameters of our integrated model enabled a quantitative description of environmentally or genetically perturbed populations. The model correctly predicted a reduction in the rate of protein synthesis in the mutant and wild-type cells grown in 0.1% glucose medium (Fig. 4H).

The loss of carbon source modulation in the TM6* and the *hxk2 hxk1* mutants may be interpreted as a collateral effect of the reduction in their growth rates, which was in the range where size is independent of λ (Fig. S3B-C) ^17,43^. However, based on their growth rate, TM6* cells grown in 2% glucose were significantly smaller than expected, similar to wild-type cells cultivated in 0.2% glucose medium (Fig. S3A-B). Furthermore, this explanation failed to address the behavior of the *snf3 rgt2 gpr1 gpa2* mutant, which is defective in extracellular glucose sensing and still exhibits nutritional modulation of cell size, albeit with alterations, despite reductions in λ like those observed in the TM6* strain (Fig. S3D, P_2%Glu/_P_EtOH_ = 1.506). Interestingly, the ethanol yield in the *snf3 rgt2 gpr1 gpa2* mutant was significantly higher than that in the TM6* and *hxk2 hxk1* mutants (Fig. S3F-H), indicating a shift in metabolism toward fermentation, which, according to the prediction of our model, seems to be a fundamental prerequisite for glucose-dependent cell size modulation (Fig. 4A). Nonetheless, in contrast to the wild-type strain, the ethanol yield values for this sensing mutant were relatively constant at all tested glucose concentrations, likely due to the impaired sugar uptake system of this mutant (Fig. S3H) ^60,61^.

Adopting the low-ATP-yield, “inefficient” fermentative metabolism instead of respiration seems a universally conserved strategy for fast-growing cells (including bacteria, fungi, and mammals), possibly reflecting a trade-off between energy yield and protein costs to optimize rapid growth under nutrient-rich conditions ^62,63^. Despite the higher ATP yield of respiration relative to fermentation, the proteome cost of energy biogenesis by respiration substantially exceeds that of fermentation and is likely the dominant cost for fast-growing cells ^62,64-68^. Therefore, forcing respiration under nutrient-rich conditions (such as in the TM6* and *hxk2 hxk1* mutants) may prevent the optimal allocation of limited proteomic resources, severely limiting the capacity of yeast cells to maximize growth and modulate their size.

Genome-wide computational models of yeast metabolism describe the salient features of yeast growth, which are simply modeled as an increase in total biomass ^69,70^. These models excel in describing steady-state situations; however, because of their nature, they cannot provide an appropriate framework for the integration of metabolism with macromolecular synthesis, cell cycle, and division. Because complex biological functions, which may involve hundreds or thousands of gene products, arise as emergent properties of a network ^71,72^ and not merely as the sum of the activities of all proteins involved in that network, dynamic molecular models are required to understand how a genotype interacts with the surrounding environment to produce a given phenotype ^73-76^.

We built the model without molecular details, because its modular structure allows for the substitution of coarse-grained components with modules that contain more molecular information. We provided a proof-of-principle validation of our model’s ability to act as a scaffold for molecularly detailed models by plugging in an existing model of the G_1_/S transition ^53^ that quantitatively describes the wild-type and mutant strains (Fig. 6). Additional cell size control mechanisms, different from those functioning in G_1_, have been reported in G_2_/M ^77,78^ and could later be introduced together with the cell cycle modules already described in the literature ^79,80^. Other high-resolution models can thus be constructed and plugged into the Metabolism and Growth modules. For instance, a recent study showed that in *Escherichia coli*, the existing mechanism for ribosomal biosynthesis regulation guarantees optimal trade-off control in resource allocation for a maximal growth rate ^81^. An analogous mechanistic model for yeast ribosomal biosynthesis may replace the current MeGro optimization step. Additionally, resource allocation optimization can be performed at the genome scale, as was recently described for *E. coli* ^82^ and applied to central metabolism in yeast, under the simplifying condition that flux and protein levels are coupled ^66,83^, as experimentally observed in *E. coli* ^84^.

In conclusion, our coarse-grained algorithm accurately recapitulates the major macrofunctions of dividing cells (metabolism, growth, and cell cycle, which require the coordinated action of thousands of gene products). This model allows us to identify the macro-function(s) that are altered by environmental or genetic perturbations. Its modular nature allows a natural way to increase the molecular details of the model. In particular, it may integrate one at a time or in combination with molecular and mechanistic models of cell subsystems, whose construction may be facilitated by sophisticated artificial intelligence approaches ^85^. Thus, the properties of mechanistically modeled functions can be analyzed within a validated framework that quantitatively describes the genetic and environmental perturbations of growing yeast populations.

### STAR*METHODS

- **KEY RESOURCE TABLE**
- **CONTACT FOR REAGENT AND RESOURCE SHARING**
- **EXPERIMENTAL MODEL AND SUBJECT DETAILS**
- **METHOD DETAILS**
  - Yeast strains and plasmids
  - Determination of growth parameters
  - Determination of DNA, RNA and protein cellular contents by flow cytometry
  - Determination of glucose consumption and ethanol production rates
  - Protein synthesis assays
  - Estimation of Oxygen Consumption Rates
  - 2-NBDG uptake assay
  - Hexokinase assay
  - Statistical analysis
- **MATHEMATICAL MODEL DETAILS**
  - *iMeGroCy* as a whole
  - The *MeGro* module
  - The optimization problem associated to a pure glucose environment
  - The optimization problem associated to a pure ethanol environment
  - Setting of the *GroCy* parameters in the *iMeGroCy* scaffold
  - Implementing asymmetrical division and genealogical age heterogeneity
  - Simulations of cell populations through *iMeGroCy*
  - Simulations of cell populations through *Hy-iMeGroCy*
  - Simulations of mutant populations
  - Sensitivity analysis
- **SOFTWARE AVAILABILITY**

### KEY RESOURCES TABLE

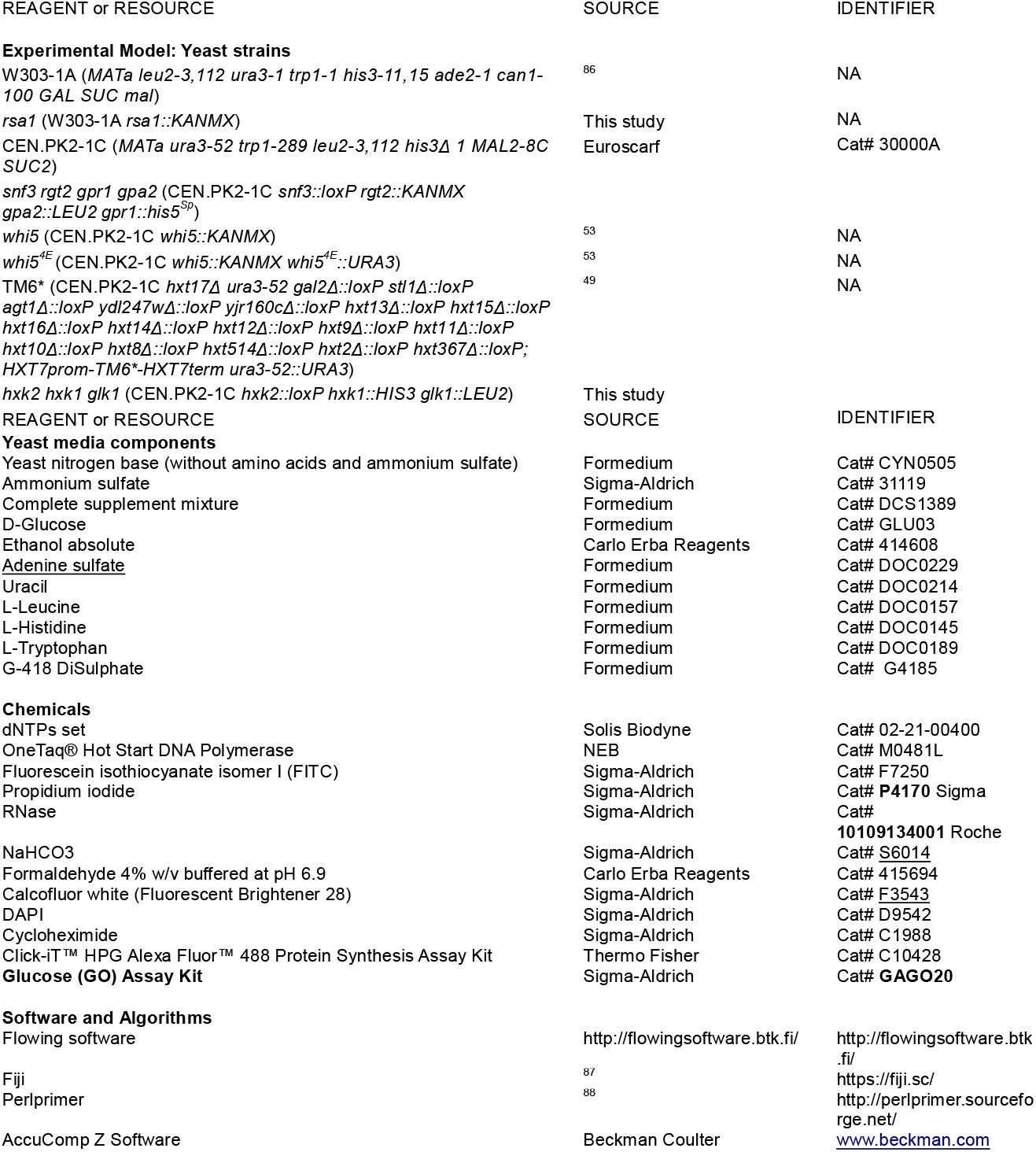

### CONTACT FOR REAGENT AND RESOURCE SHARING

Further information and requests for resources, reagents and modeling should be directed to and will be fulfilled by the Lead Contact, Marco Vanoni.

### EXPERIMENTAL MODEL AND SUBJECT DETAILS

All strains used in this study were derivatives of *S. cerevisiae* W303-1A ^86^ and CEN.PK2-1C (Euroscarf Cat# 30000A). The Key Resources Table reports a complete list of the strains used in this study.

## METHOD DETAILS

### Yeast strains and plasmids

Cassettes for *WHI5* and *RSA1* gene deletions were generated via PCR by using genomic DNA of specific haploid strains from the deletion collection (Euroscarf) as template and short primers (∼20bp long, sequence available upon request) designed at least 100 bp upstream/downstream of the ATG and stop codon using the Perlprimer software (http://perlprimer.sourceforge.net/ ^88^), as described ^89^. Alternatively, we generated deletion mutants by the PCR-mediated gene disruption method (short flanking homology *loxP::marker::loxP/Cre* recombinase system ^90^. Cells were transformed by the standard lithium acetate procedure ^91^. Deletion of the targeted gene sequences in transformants was confirmed by colony PCR. Whenever needed, the markers used for gene disruptions were removed by inducing the recombination of the flanking *loxP* sequences through the expression of the Cre recombinase ^90^. For inactivation of *HXK1*, an *hxk1::HIS3* disruption cassette was amplified by PCR using genomic DNA from a *hxk1*-null strain as a template ^92^. *GLK1* was disrupted using a *glk1::LEU2* cassette obtained by digesting the PWA40 plasmid with *NcoI/SacI* ^92^.

*gpa2* null strains were constructed by one-step gene replacement using a *gpa2::LEU2* disruption cassette obtained by digestion with *PstI* of the pUC19-*gpa2::LEU2* plasmid ^93^.

A DNA fragment encoding the *whi54E* mutant was synthesized *de novo* by Eurofins (www.eurofins.com) and subcloned into the YIplac211 integrative plasmid^66^ under the *WHI5-545*_*pr*_ native promoter^42^, yielding the construct YIplac211-*545*_*pr*_*-WHI54E* ^53^. Single-copy genomic integration of the construct at the *URA3* locus was verified by quantitative PCR as described ^53^.

### Determination of growth parameters

Yeast cultures were grown in synthetic complete minimal medium, containing 0.67% (w/v) yeast nitrogen base (YNB), appropriate quantities of the ‘drop-out’ amino acid–nucleotide mixture and supplemented with either 2% (w/v) glucose, 0.05% (w/v) glucose or 2% (v/v) ethanol.

The growth of cultures was monitored as increase in cell number using a Coulter Counter model Z2 (Beckman Coulter). Cell size analysis was performed using a Coulter Z2 Particle Cell Analyzer (Beckman-Coulter). The fraction of budded cells (FB) was scored by direct microscopic observation on at least 400 cells, fixed in 3.6% formaldehyde and mildly sonicated.

The fraction of parent budded (FPP) and unbudded (FPNB), daughters budded (FBD) and unbudded (FDNB) cells inside the population was determined by bud scar analysis after 10 min staining with Calcofluor white (25 μM in PBS, protected from light): at least 800 cells were scored by direct microscopic counting under a Nikon Eclipse E600 fluorescence microscope, equipped with a 100X, 1.4 oil Plan-Apochromat objective, and standard DAPI filter set. Images were digitally acquired using a Leica DC 350F camera and processed with the FiJi software (https://fiji.sc/ ^87^).

The formulas used to calculate the length of the budded period (T_B_), of the binucleated phase (T_G1*_), the average parent cycle time (T_P_) and the average daughter cycle time (T_D_) were ^35^

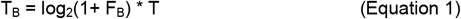

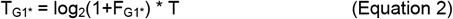

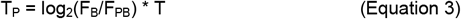

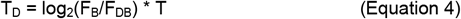

where T is the overall duplication time of the population, F_B_ is the percentage of budded cells, and F_PB_ and F_DB_ are the fractions of budded parents and daughters in the whole population, as determined by bud scar analysis. Due to the asymmetrical division, the parent and the daughter cycle times must satisfy the equation

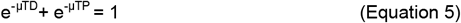

where μ is the growth rate given by

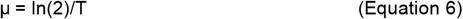

### Determination of DNA, RNA and protein cellular contents by flow cytometry

Samples of growing cultures (at least 2 * 10^7^ cells) were collected and fixed in 70% ethanol before the cytofluorimetric analysis. For protein staining, cells were washed once with cold PBS (3.3 mM NaH_2_PO_4_, 6.7 mM Na_2_HPO_4_, 127 mM NaCl. 0.2 mM EDTA, pH 7.2), resuspended in 1 ml of freshly prepared protein staining solution (fluorescein isothiocyanate (FITC) 50 µg/ml in 0.5 M NaHCO_3_) and incubated for 1 h in ice protected from light. After incubation cells were washed three times with PBS and resuspended in 1 ml PBS. For DNA staining, cells were washed once with PBS, resuspended in 1 ml of PBS with RNAse 1 mg/ml and incubated overnight at 37°C. Cells were then washed once with cold PBS resuspended in 1 mL of propidium iodide staining solution (0.046 mM propidium iodide in 0.05 M Tris–HCl, pH = 7.7; 15 mM MgCl_2_) and incubated for 30 min in and in the dark. The RNAse treatment was omitted for RNA staining.

For DNA/Protein bi-parametric analysis, RNA-treated cells were resuspended in 1 mL of a 1000-fold diluted FITC staining solution (50 ng/mL FITC in 0.5 M NaHCO_3_) and incubated for 30 min in ice protected from light; cells were then washed three times with PBS and resuspended in DNA staining solution ^87^. Cell suspensions were sonicated for 30 s before the analysis, which was performed with a FACSCalibur (Becton Dickinson) instrument equipped with an Ion-Argon laser at 488 nm laser emission. The sample flow rate during analysis did not exceed 500-600 cells/s. Typically, 30,000 cells were analysed for each sample.

The average protein content (P) of the entire population, at the beginning of the cell cycle (P_0_) and the onset of DNA replication (P_s_) were determined as average fluorescence intensity of appropriately gated cells from the density plot derived from FACS analysis of double DNA/protein stained cells. Plot generation, data analysis and gating process were performed with Flowing software (http://flowingsoftware.btk.fi/). P_s_ and P_m_ (the protein content at cellular division) are correlated population parameters: their ratio h is given by the equation ^17^

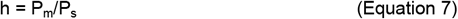

The value of P_P_ (the average protein content of parent cells) can be calculated from either ^17^

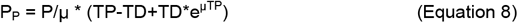

Or

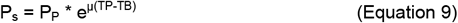

### Determination of glucose consumption and ethanol production rates

At different time points samples of culture were collected and centrifugated. Extracellular levels of glucose and ethanol in the supernatants were evaluated by ^1^H-NMR as previously reported ^94^. Alternatively, glucose consumption and ethanol production were evaluated through a Hyperlab automatic multi-parametric analyzer SMART (Steroglass, San Martino in Campo, Perugia, Italy) and specific enzymatic kits for ethanol and glucose detection (Steroglass). Hyperlab analyzer, managed by the Hyperlab SMART Software, handles samples, reagents and dilutions in a reproducible way without the need of human processing. Metabolite concentration (g/L) were automatically calculated with appropriate algorithms from measured absorbance values by using standard calibration curve as reference.

### Protein synthesis assays

We quantified Protein synthesis rate with the “Click-iT HPG Alexa Fluor 488 Protein Synthesis Assay Kit” (Thermo Fisher Scientific), with minor adaptations from the manufacturer’s protocol. The Kit detects the incorporation into nascent proteins of HPG (L-homopropargylglycine), a methionine analog containing an alkyne moiety, through labeling with a fluorescent azide (Click-iT reaction). Yeast cells were grown overnight at 30 °C in SC methionine-free medium. For each sample, 1*107 cells were collected and resuspendend in 100 μl of growth medium supplemented with 50 μM HPG. After 15 min incubation at 25° C, cells were washed once with ice-cold PBS and fixed for 15 minutes at room temperature with 100 μl of 3.7% formaldehyde in PBS. After fixation, cells were washed twice with 1 ml of 3% BSA in PBS, resuspended in 100 μl of permeabilization buffer (0.5% Triton X-100 in PBS) and incubated for 20 min at room temperature. Permeabilized cells were then washed twice with 1 ml of 3% BSA in PBS, and the “Click-iT reaction” was performed: cells were resuspended in 100 μl of Click-iT reaction cocktail (freshly prepared according to manufacturer’s datasheet) and incubated for 30 minutes at room temperature, protected from light. After removal of the reaction cocktail, cells were washed once with 100 μl of rinse buffer and once with 1 ml of PBS. Cells were finally resuspended in 1 ml PBS and mildly sonicated (twice for 15 s) before analysis. The fluorescence arising from incorporated HPG-Alexa Fluor 488 was detected by flow cytometry with a FACScalibur instrument (Becton Dickinson). The mean fluorescence intensity for each strain/growth condition was evaluated from 50000 cells and normalized to the value of the wild type strain. Samples incubated in the absence of HPG were used as negative controls. If required, protein synthesis was inhibited by pre-incubating cells for 60 min with 100 μg/ml cycloheximide before HPG addition.

### Estimation of Oxygen Consumption Rates

Oxygen consumption in intact cells was measured at 30° C using a “Clark-type” oxygen electrode in a thermostatically controlled chamber (Oxygraph System, Hansatech Instruments), essentially as described ^17^. Briefly, 2 mL of cell suspension at a concentration of ∼5×106/mL were quickly transferred from flask to the oxygraph chamber. The basal/routine oxygen consumption rate (*J*_*R*_) was determined from the slope of a plot of O_2_ concentration against time, divided by the cell number. Treatment with 37.5 mM Triethyltin bromide (TET, a lipophilic F0F1-ATPase inhibitor; Sigma) allowed the determination of the non-phosphorylating respiration rate due to proton leakage (*J*_*TET*_), whereas the maximal (uncoupled) oxygen consumption rate (*J*_*MAX*_) was measured after addition of the protonophore Carbonyl cyanide-*4*-(trifluoromethoxy)phenylhydrazone (FCCP, Sigma) at 10 µM and of saturating amounts of ethanol (100 mM) as respiratory substrate. The addition of 2 mM antimycin A (Sigma) accounted for non-mitochondrial oxygen consumption. All assays were performed at least in biological triplicate. The net respiration (netR) was estimated by subtracting *J*_*TET*_ from *J*_*R*_ and used to calculate the net routine control ratio *netR/J*_*MAX*_ ^35^.

### 2-NBDG uptake assay

Glucose uptake rate was estimated by using the fluorescent non-hydrolysable analogue 2-NBDG (2-Deoxy-2-[(7-nitro-2,1,3-benzoxadiazol-4-yl)amino]-D-glucose; Sigma). Exponential phase cells were harvested by centrifugation, washed once with fresh medium (w/o glucose) and resuspended at 1*108 cells/ml in fresh medium (w/o glucose). 2-NBDG (25 mM stock solution in DMSO) was added to 50 uL of cell suspension (final concentration 0.2-0.4 mM). Cells were incubated for 30 min at 30 °C, collected by centrifugation, washed twice with ice cold PBS and resuspended in 1 mL PBS. After mild sonication (2 x 15 s) the mean incorporated fluorescence intensity was detected by a FACScalibur instrument (FL1 filter) from 30000 cells and normalized to the value of the wild type strain grown in 2% glucose medium. Samples incubated without 2-NBDG were used as negative controls.

### Hexokinase assay

Hexokinase activity was measured essentially as described ^17^, with minor modifications. Briefly, ∼3*108 exponentially growing cells were harvested by centrifugation and washed once in ice-cold distilled water. Cells were disrupted by glass bead lysis in 40 mM MES-Tris buffer (pH 6.8) containing 2 mM PMSF. Protein concentration of samples was measured by Bradford method and 200 mg of crude extract were used for the phosphorylation assay. Samples were supplemented with 20 mM D-glucose, 10 mM ATP, and 5 mM MgCl_2_ and incubated at 30°C. The amount of glucose remaining at each time point was measured by using a glucose-oxidase-peroxidase reaction kit (Sigma). Glucose phosphorylation activity was evaluated from the total amount of glucose consumed per unit of time.

### Statistical analysis

Data are reported as means□± □SDs from at least three independent experiments. Statistical significance (indicated with “*”) of the measured differences was assessed by two-sided Student’s t-test (p□ < □0.05).

## MATHEMATICAL MODEL DETAILS

The mathematical model of the budding yeast *S. cerevisiae* deals with its main cellular activities, namely metabolism, growth and cycle. The model is denoted in the following as *integrated-MeGroCy* model (*iMeGroCy*, shortly). A preliminary version of the *iMeGroCy* model, limited to single cell and accounting only for the external glucose as cell nutrient, can be found in ^17^. Here, instead, *iMeGroCy* is extended to cell populations, allowing the analysis of protein distributions, while the metabolism module is updated in order to allow cell growth in the presence of ethanol as cell nutrient.

### iMeGroCy as a whole

The model is conceived to be modular and hierarchical. The growth activity combined to the other two main cellular activities of metabolism (*MeGro*) and cycle (*GroCy*) define the modular building blocks constituting the coarse-grained backbone of *iMeGroCy* (see Fig. 2A). Both modules may, in principle, be substituted (in part or totally) by a finer (possibly molecular) blow-up. The wiring among the modular building blocks is functional to the correct setting and working modes of the integrated model as a whole. A functional scheme is reported in Fig. 2B, where *MeGro* is enlisted to sense the environmental nutrient (e.g., the external glucose or the external ethanol) and to process the ethanol yield information coming from experimental data in order to set the exponential growth rate λ and the ribosome-over-protein ratio ρ coming at steady-state as outputs of an optimization algorithm aiming at maximizing the growth rate. Unlike *MeGro, GroCy* is a dynamical model. The *MeGro* outputs λ and ρ enter *GroCy* as inputs, allowing to set the ribosome and protein dynamics parameters, constituting the growth module. Growth and cycle are linked together through a molecular trigger representing the Cln3/Far1 dynamical interplay. The growth-to-trigger link is rendered by constraining the total amount of Cln3 to the overall protein content. The cycle module is composed of three consecutive timers, T_1_, T_2_, and T_B_. T_1_ and T_2_ are named after the definition of ^17^ and their union provides the G_1_ phase: the former refers to the period a newborn cell takes to activate the G_1_/S regulon and is formally measured by the time the regulon inhibitor Whi5 takes to exit the nucleus; the latter refers to the time cyclins Clb5/6 (responsible for the onset of the S phase) take to get rid of their inhibitor Sic1. The trigger-to-cycle link is rendered by the fact that the onset of the final part of T_1_ is regulated by the molecular trigger and occurs when the upstream cyclin Cln3 gets rid of its inhibitor Far1. T_B_ includes the rest of the cycle, namely, phases S, G_2_, and M. Part of the *GroCy* parameters are influenced by and vary according to the nutrient environment. The last 20% of timer T_B_ refers to the G_1_* phase, according to which 2 nuclei are present in a yet single cell. A RESET function implements nuclear division. Further details concerning the *iMeGroCy* for single cells can be found in ^17^.

### The MeGro module

The present version of *MeGro* is basically an extension of the model reported by ^17^, that was derived from the “self-replicator” model proposed by ^17^. Both (basic and extended) versions of *MeGro* are conceived to highlight the common patterns connecting growth rate-dependent regulation of cell size, ribosomal content and metabolic efficiency in yeast Saccharomyces cerevisiae, showing how such patterns arise from the growth rate maximization. The extended version of *MeGro* allows the metabolic engine to work consuming both external glucose and ethanol (the basic model takes into account the external glucose as the only nutrient of the cell ^17^), so providing the growth condition of the cell when glucose or ethanol are present in the culture. See ^17^ for more details on the formulation of the basic model in glucose environment.

*MeGro* accounts for five classes of protein-like players having enzymatic activity, and seven kinds of metabolites. The five players with enzymatic activity (square blocks in the *MeGro* scheme of Figure. 2A) are (i) the hexose transporters, *‘hxt’*, (ii) the glycolytic enzymes, *‘gly’*, (iii) the ribosomes, *‘rib’*, the enzymes of (iv) respiration and (v) fermentation pathways, *‘resp’* and *‘ferm’* respectively. Five kinds of metabolites are involved in metabolic conversions (green ovals in the *MeGro* scheme of Fig. 2A), namely the (a) extracellular and (b) intracellular glucose, *‘glc*_*ex*_*’*, and *‘glc*_*in*_*’* respectively, (c) extracellular and (d) intracellular ethanol, *EtOH*_*ex*_ and *EtOH*_*in*_, respectively, and (e) pyruvate, *‘pyr’*; other two kinds of metabolites are involved in energy production/consumption: (d) ATP and (e) ADP. In the following we will indicate by *c*_*x*_, *x ∈ {hxt, gly, rib, resp, ferm}/{glc*_*ex*_, *glc*_*in*_, *EtOH*_*ex*_, *EtOH*_*in*_, *pyr, ATP, ADP}*, the protein/metabolite player concentrations, (mM), and by *ν*_*x*_, *x ∈ {hxt, gly, rib, resp, ferm}*, the metabolite fluxes, (mM/h) catalysed by a specific protein-like player *x*.

By looking at the *MeGro* scheme depicted in Fig. 2A, but also at its equations reported in the following, we note that the model accounts for two respiratory pathways catalysed by the same class of proteins, *c*_*resp*_ (represented by the red square “resp” in Fig. 2A), but fed by two different metabolites, i.e. pyruvate, *c*_*pyr*_, and ethanol,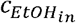. The mentioned respiratory pathways produce two distinct fluxes, i.e. *ν*_*resp*_ and *ν*_*resp2*_. Moreover, two opposite pathways concerning glucose metabolism are taken into account by the present model formulation: (i) the glycolysis, converting glucose into pyruvate, represented by the flux *ν*_*gly*_; (ii) the gluconeogenesis, producing glucose from ethanol, modelled by the flux *ν*_*gly2*_. Both fluxes are catalysed by the same class of glycolytic enzymes, that is *c*_*gly*_ (represented by the red square “gly” in Fig. 2A). The inclusion of the gluconeogenesis pathway in the present model formulation allows *MeGro* to sustain the synthesis of the building blocks of the cell by two distinct pathways, one fed by the external glucose and the other one by the external ethanol.

Going through the details of the model equations, the protein synthesis is modeled as a proper fraction α_*x*_ of the total ribosomal flux *ν*_*rib*_, thus, in exponential growth conditions we have the following steady-state constraints (see ^17, 17^ for more details):

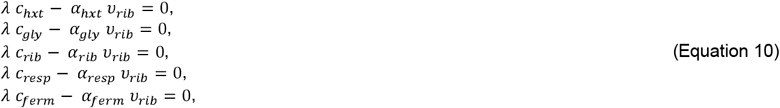

where λ, (h^-1^), is the specific growth rate and

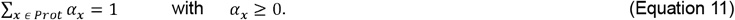

External glucose and external ethanol are two model inputs. Assuming a growth medium with an infinite volume, the external concentrations of glucose and ethanol, 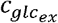 and 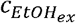, can be consequently considered constant. Conversely, the dynamics of the internal metabolites 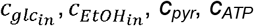 are determined by the combination of the fluxes of synthesis and degradation:

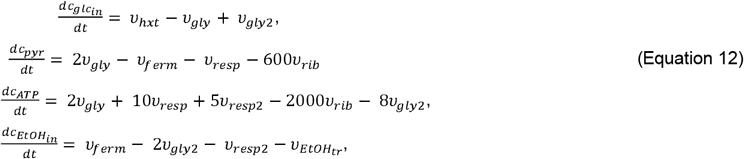

written according to the following chemical reaction scheme:

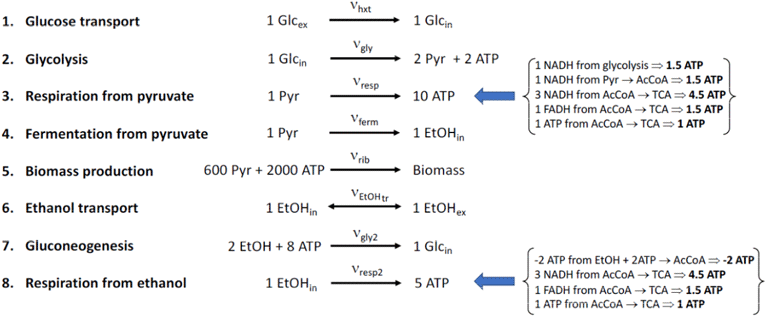

At steady-state the following constraints on the metabolic fluxes can be written:

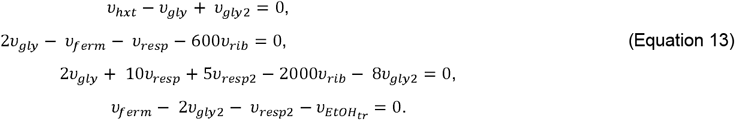

The total amount of *ADP* and *ATP* is constant, according to the following relationship

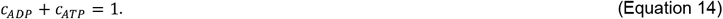

All protein/metabolite concentrations *c*_*x*_ are such that *c*_*x*_ ≥ 0.

Concerning the kinetic laws of the metabolic fluxes, they are modelled using the Michaelis-Menten formalism:

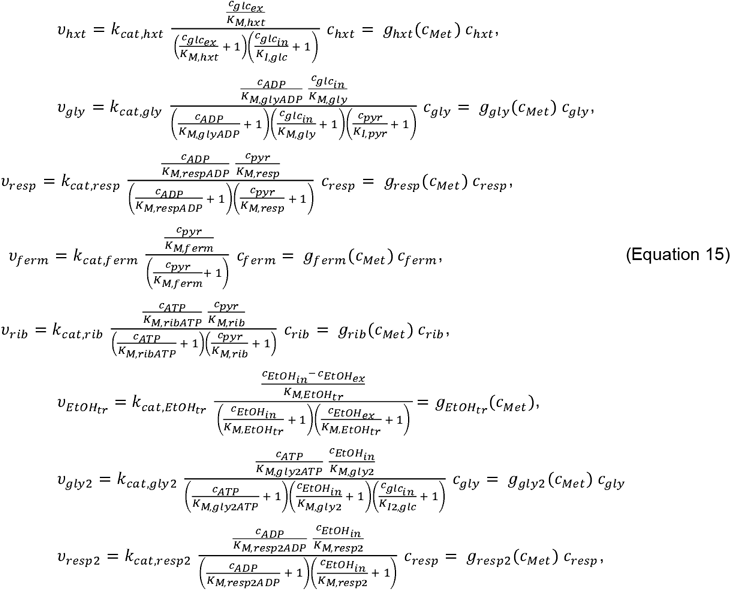

where *c*_*Met*_ is the vector of the seven metabolite concentrations. Equations from (10) to (15) define the set of algebraic-differential equations of *MeGro*. The exponential growth rate λ can be maximized as a function of the external glucose concentration 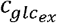, or of the external ethanol concentration, 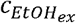, with the fractions α_*x*_ as optimization variables, and subject to exponential growth constraints, equations (10), flux balance constraints, equations (13), Michaelis-Menten flux equations, equations (15), and feasible constraints, equations (11), (14).

The values of *MeGro* input parameters are reported in Table S2, with the tuning of *MeGro* mostly discussed in ^17^. Regards to the new parameters, they have been set in order to ensure that the new version of *MeGro* is consistent with the old version ^17^ in glucose environment. More in details, the parameters of 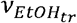 were chosen in order to make the exit transport of 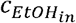more favorable than the recycling throughout *ν*_*gly2*_ and *ν*_*resp2*_ in the absence of external ethanol (i.e. when 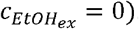 . Indeed, we fixed the parameters 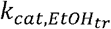and 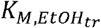 to a very high (about 150 times *k*_*cat,ferm*_) and, respectively, to a low value (about 100 times lower than *k*_*M,ferm*_). So, when glucose is the only nutrient, it is *v*_*gly*2_ ≪*v*_*gly*_and *v*_*resp*2_≪ *v*_*resp*_, that is 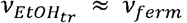. Finally, experimental data on MDT of WT strain CEN.PK in ethanol environment (2%) have been used to fine tuning the kinetic parameters of the new branches of *MeGro* (in particular *v*_*gly*2_ and *v*_*resp2*_).

The optimal value of λ together with the fractions α_*x*_, the protein/metabolite concentrations and the protein fluxes provide a first level of *MeGro* outputs. Additional outcomes, computed by properly exploiting concentrations and fluxes, are (i) the *fermentative ratio F*,

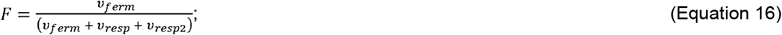

(ii) the *ribosome-over-protein ratio* ρ, given by the following expression

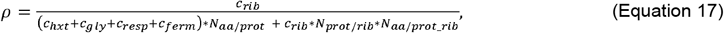

where *N*_*prot/rib*_ is the average number of ribosomal proteins per ribosome, *N*_*aa/prot-rib*_ is the average number of amino acids per ribosomal protein, and *N*_*aa/prot*_ is the average number of amino acids per generic protein; so the expression (17) returns the *ribosome-over-protein ratio* in terms of number of ribosomes over the total amount of polymerized amino acids within the five macro-classes of synthetized players. Giving the total amount of proteins by means of the total number of polymerized amino acids allows *MeGro* to straightforwardly communicate with *GroCy* (see Fig. 2A), which accounts for cell size in terms of number of polymerized amino acids too;
(iii) the yield of ethanol *Y*_*EtOH/glc*_ produced in the presence of glucose nutrient:

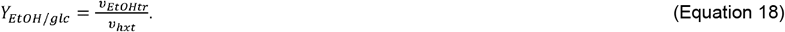
(iv) the protein investments

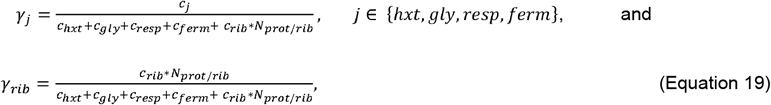

which provide the percentage of proteins that the cell produces within each class of players having enzymatic activity.

### The optimization problem associated to a pure glucose environment

As far as the numerical implementation of the optimization problem is concerned, disregarding the equations of the additional model outputs (17-19), the problem counts: (i) 25 unknown variables, i.e. the 5 concentrations of the protein players, 5 metabolite concentrations (considering only the unknown metabolites, since 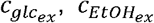 are fixed based on the external environment), the 8 fluxes, the 5 fractions of ribosome flux *α j* and *F*; (ii) 20 algebraic constraints, i.e. Eqs. (10-11) and (13-15), including also the equation of *F* (16). So, from the theoretical point of view, the optimization problem has 5 degrees of freedom.

By choosing 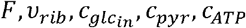 as independent variables, from Eqs. (10-11) we obtain the explicit expression of the growth rate

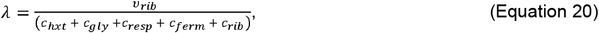

as well as the equations *α* _*j*_ =*C*_*j*_*/(C*_*hxt*_ *+ C*_*gly*_ *+C*_*resp*_ *+C*_*ferm*_ *+C*_*rib*_*), j* ∈ {*hxt, gly, resp, ferm, rib*} allowing to derive *α* _*j*_ from the knowledge of the concentrations *C* _*j*_.

From the Michaelis-Menten flux equations (15), we get the relations

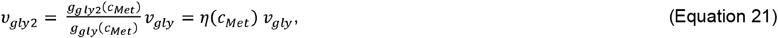

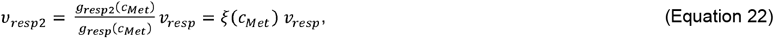

where the two ratios *η* (*CMet), ξ* (*CMet)*are explicit functions of the metabolite concentrations only. By properly exploiting the flux balance constraints (13) and the fermentative ratio definition (16), we can express all the fluxes in terms of *F*, the ribosomal flux *v*_*rib*_ and the metabolite concentrations as follows:

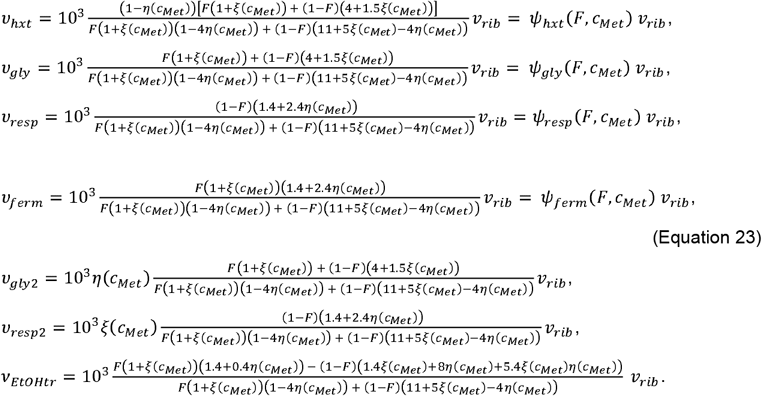

By suitably inverting Eqs. (15) w.r.t. *C* _*j*_ and taking into account Eqs. (23), we get the following explicit expressions for the concentrations of the protein-like players:

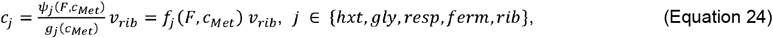

where *ψ* _*rib*_ is formally set equal to 1. From Eqs. (24) and (20), we get

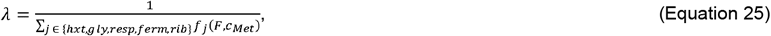

where the explicit dependence on *v*_*rib*_ disappears. Among the entries of the vector 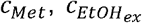 and 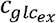 are fixed, *C*_*ADP*_ is univocally determined by *C*_*ATP*_ from Eq. (14), while 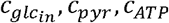, are free variables moved by the optimization routine together with *F*, with *v*_*rib*_ and 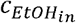 still involved by the constraints in Eqs. (15), (23).

However, we verified by simulation that is “practically” independent of *v*_*rib*_ in glucose environment (i.e. 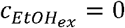) so that the actual degrees of freedom of the optimization problem is 4.This fact is somehow implied by the tuning of the model parameters achieved in order to have *v*_*gly*2_ ≪*v*_*gly*_and *v*_*resp*2_≪ *v*_*resp*_ when 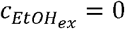, that means, in order to make the transport of ethanol more favorable than its recycling in glucose environment. This property has been numerically verified by preliminary simulations performed for 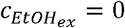 and 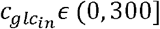 (with *F ϵ* [0,1] and 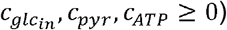), revealing that *η* (*CMet), ξ* (*CMet)* assume always low values, at most of the order of magnitude of 10^-3^. This numerical property implies that *ψ*_*j*_ (*F,C*_*Met*_) ≈*ψ*_*j*_ (see equations (23)) and then *F* _*j*_ =(*F,C*_*Met*_)*= ψ*_*j*_ *(F)/ g*_*j*_ (*C*_*Met*_*), j* ∈ {*hxt, gly, resp, ferm, rib*}. Therefore, since *g*_*j*_ (*C*_*Met*_), *j* ∈ {*hxt, gly, resp, ferm, rib*}, are not dependent on 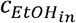, we can conclude that is “practically” independent of *v*_*rib*_ and then the numerical optimization can be performed w.r.t. the four free variables 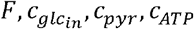 (fixing *v*_*rib*_ to an arbitrary value, for instance 1).

Similarly to the old version presented in ^17^, *MeGro* can treat the fermentative ratio *F* also as an input (so reducing the number of free variables to 3), rather than an output, when no ethanol is provided by the experimenter, thus allowing the modeler to compute the optimal growth rate (as well as all the other model outputs) according to the fixed glucose and different values of *F*. See the paper ^42^ for more details on *MeGro* implementation.

In order to perform data-driven numerical experiments, it is very useful to derive a tool that allows to translate experimental data into model inputs. In particular, from the model equations given above, it is possible to derive an algebraic constraint which relates the fermentative ratio *F* to the yield of ethanol *Y*_*EtoH/glc*_. More in details, according to the ethanol yield definition (18) and to the expressions of *v*_*EtoHtr*_ and*v*_*hxt*_ given by Eqs. (23), we get

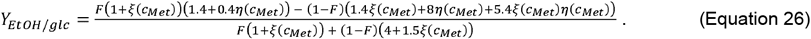

Since, with the given parameter setting, it is η(c_*Met*_), ξ ;(c_*Met*_)≪ 1 when 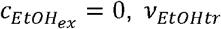 and*v*_*hxt*_ in Eqs. (23), can be reasonably approximated as:

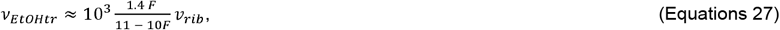

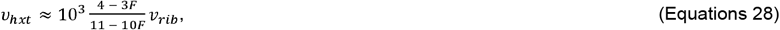

providing the following approximated expression for the ethanol yield:

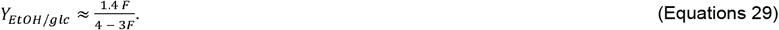

Eq. (29) is bijective, thus it can be inverted, providing the fermentative ratio *F* associated to a given ethanol yield:

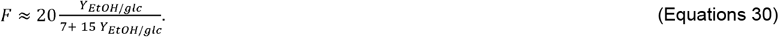

This last equation will be exploited to feed *MeGro* with the fermentative ratio associated to experimentally measured values of the ethanol yield (see Fig. 2B).

### The optimization problem associated to a pure ethanol environment

When 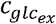 vanishes, from the kinetic equation (15) it is *v*_*hxt*_ = 0 and *v*_*gly2*_= *v*_*gly*_, that means η (c_*Met*_)=1 for any value of c_*Met*_ . Since the condition *v*_*hxt*_ = 0 is satisfied for any value of *v*_*hxt*_ = 0, the optimization routine would bring *c*_*hxt*_ (as well as *α*_*hxt*_) to zero for maximizing λ(see Eq. (20)). However, in order to perform more sound and realistic numerical experiments it is possible to fix the investment *γ*_*htx*_ on the transporter ‘hxt’ (see Eq. (19)) on the basis of experimental data. Therefore, adopting a data driven approach, we can set *y*_*htx*_ to the experimental percentage of proteins that the cell allocates for the class ‘hxt’, so inferring the corresponding concentration from the data as

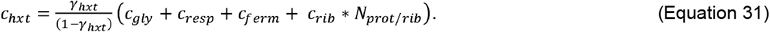

Moreover, in order to simulate the growth in ethanol condition it is reasonable to set F = 0 that implies *v*_*ferm*_ = *c*_*ferm*_ = *α*_*ferm*_ = *γ*_*ferm*_ = 0. Note that, such a condition is the best option that the optimization could set, even if *F* were not fixed. Indeed, preliminary simulations showed that γ increases for decreasing (fixed) values of *F* (while for high values of *F*, i.e. approximately *F* > 0.4, no feasible solution is found).

From Eqs. (20) and (31), and taking into account *F* = 0, we get the following expression for the growth rate

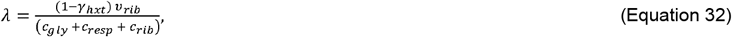

Where 0 ≤ *γ*_*hxt*_ ≤1. Moreover, from Eqs. (23-24), we obtain

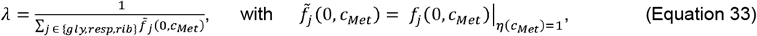

where the explicit dependence on *v*_*rib*_ disappears again. However, from a theoretical point of view, the value of *v*_*rib*_ could still have an indirect influence on the growth rate, since it influences the value of 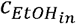. Conversely, from preliminary numerical simulations, it turns out that *v*_*rib*_ is not practically identifiable and then it has been arbitrarily fixed to 1 as in the glucose case. Note that the chosen value of *v*_*rib*_ actually influences the absolute value of 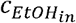, but it has no practical influence on the main MeGro outputs, *λ, ρ* (which are fractional quantities, not dependent on the absolute values of the other model variables, as explained above).

### Setting of the *GroCy* parameters in the *iMeGroCy* scaffold

The following procedure concerns the setting of the model parameters for an average single cell in different nutritional conditions. The idea is to mimic the cellular organization as flow of information: different nutritional environments (experimental data) stimulate different metabolic responses (*MeGro* outputs), and these eventually modulate downstream growth and cycle processes (*GroCy* setting).

Experimental data are generated on 6 batch cultures with different glucose concentrations and for ethanol as nutrient. The *MeGro* parameters ^17^ have been fixed equal for all the nutritional environments. The experimental data (Tables S1A-C) are:

1. the ethanol, *c*_*EtOH_ex*_, and glucose concentration, *c*_*glc_ex*_, this last ranging 5%, 2%, 0.5%, 0.2%, 0.1%, 0.05%;
2. the Mass Duplication Time, MDT, from which the exponential growth λ is readily computed as: λ = log(2)/MDT;
3. the yield of ethanol, Y_EtOH/glc_, for glucose concentrations.

These experimental data are integrated with the fermentative ratio *F* (i.e. the fraction of fermentative flux provided for different glucose concentrations), that *MeGro* univocally associates to the ethanol yield (like in ^17^), thus providing a bijection from the external glucose concentration to *F*. In order to utilize a smoother 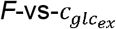 relationship, we used the best-fitting saturating function

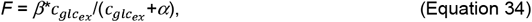

where the values of *α* and *β* have been estimated by minimizing the mean square error with respect to the available experimental data. The estimated values of the above parameters are *α* = 0.6059 and *β* = 0.9790 and the fitting curve is depicted by the blue solid line of Fig. 1D, where the squares correspond to experimental data. The smoothed values of F will be exploited instead of the raw experimental ones, in order to filter out measurement noises. In case of ethanol nutrient, *F* = 0.

The fermentative ratio *F* feeds *MeGro* as an input, together with the glucose concentration (see Fig. 2B for the block diagram scheme describing the whole procedure of *iMeGroCy* parameter setting). In turn, *MeGro* solves an optimization algorithm providing the maximum growth rate λ (and the related ribosome-over-protein ratio ρ) according to the inputs above and to the model constraints that involve fluxes and metabolites concentrations. *MeGro* outputs help to set also some of the growth parameters of *GroCy*. Indeed, the mathematical analysis carried out on the model growth equations (^53^; ^53^ has shown that, according to an approximation suggested by feasible model parameters, the balanced exponential growth condition provides the following relationship

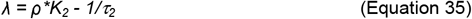

where *K*_*2*_ and *τ*_*2*_ refer to the ribosome production rate per number of polymerized amino acids and the ribosome degradation time constant, respectively. As a matter of fact, given the pair (λ,ρ), associated to a given glucose environment, eq.(35) provides a manifold onto which parameters (*K*_***2***_, *τ*_***2***_) can be chosen. The values of (ρ, *K*_***2***_), related to the optimal growth rate, are reported in Table S3 as inputs of the *GroCy* module.

We set other growth parameters (Table S3) accordingly to ^53^ and ^17^, regardless of the ethanol/glucose concentration. These growth parameter values are in general agreement with experimental data summarized in ^53^.

For the molecular trigger mechanism, most parameters do not vary with the glucose concentration. Consistently with experimental findings ^53^, the initial amount of *Far1* - which is set equal to 240 molecules for glucose concentrations from 0.1% up to 5% - is reduced to 170 molecules at 0.05% glucose concentration and to 100 molecules for ethanol. The triggering mechanism accounts for the setting of the first part of timer *T*_*1*_, namely *T*_*1a*_ (see ^17^ for the details).

The second part of timer *T*_*1*_, namely *T*_*1b*_, is strongly related to the cell size and is set according to the following constraint

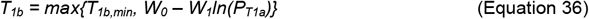

*T*_*1b*_ *= max{T*_*1b,min*_, *W*_*0*_ *– W*_*1*_*ln(P*_*T1a*_*)}* (Equation 36) where *T*_*1b, min*_ is the minimum length of *T*_*1b*_ and *P*_*T1a*_ is the cell size at the end of *T*_*1a*_. *T*_*1b, min*_ has been arbitrarily fixed at 1 min for all glucose concentrations, while *W*_*0*_ and *W*_*1*_ are chosen to ensure an appropriate *T*_*1*_ length and a good fit with the experimental data. To this end, they have been fixed equal for 5 out of 6 glucose concentrations, allowing them to vary only for the poorest mean, 0.05% glucose concentration. This choice is motivated by the fact that temporal parameters significantly vary only for the lower glucose concentrations, (e.g. 0.05% glucose, Table S2A).

We fixed Timer *T*_*2*_ length at 10 min for Daughters and Parents at any glucose concentration, a value that falls within the range of variability reported in ^53^. Conversely, timer *T*_*B*_ slightly varies according to the glucose concentration. Experimental findings show that the budded phase is kept roughly constant for higher glucose concentrations (*T*_*B*_ = 83 min, 81 min, 82 min for 5%, 2% and 0.5% glucose respectively), and slightly increases for lower glucose concentrations (*T*_*B*_ = 87 min, 87 min, 92 min for 0.2%, 0.1% and 0.05% glucose respectively). In our simulations we allowed the same general trend with two distinct values for *T*_*B*_ (a lower one for glucose 5%, 2%, 0.5%, and a slightly higher one for glucose 0.2%, 0.1%, 0.05%). These values vary for the ethanol environment, allowing longer time periods. Table S3 summarizes the input values of the parameters *T*_*1b,min*,_ T_2_, T_B_ (min), *W*_*0*_, *W*_*1*_, used in the numerical simulations.

### Implementing asymmetrical division and genealogical age heterogeneity

When a yeast cell buds, a chitin ring, called bud scar, builds up at the bud isthmus and remains on the parent cell after the bud has separated (reviewed in ^53^). Bud scars on intact cells can be visualized by fluorescent dyes (Calcofluor, Primulin) in fluorescence microscopy. Since each new bud starts at a new site, it is possible to determine the number of bud scars present on the surface of a parent cell and consequently to establish the genealogical age ‘*k’* of the parent cell, meaning the age of the parent cell equal to the number of daughters it has generated (i.e., *k = s*). So, denoting by “*P*_*k*_” a parent cell of age ‘*k’*, a cell *P*_*1*_ has one bud scar since it has completed a cycle, a cell *P*_*2*_ has two bud scars since it has completed two cycles, and so on. On the other hand, a cell without bud scars (*s* = 0) is a daughter cell and it has not yet completed a cycle. *iMeGroCy*, however, distinguishes the genealogical age of the daughter cells: it can be 1 if the daughter is born from another daughter, while it is *k > 1* if the daughter is born from a parent *P*_*k-1*_. We denote by “*D*_*k*_” a daughter of genealogical age ‘*k’*.

At division each parent receives the mass it had at budding, whereas the mass synthesized during the budding phase goes to the newborn daughter. It follows that in parents, cell mass at budding increases with genealogical age. Experimental evidence shows that the higher is the genealogical age (i.e., the number of bud scars), the smaller is the increase in size at budding from one generation to the other (reviewed in ^53^). The reduction with genealogical age of the cell size increase has been explained by mechanical stress of the cell wall, which increases with cell size ^53^; ^53^. To account for the aforementioned behavior, when dealing with a parent cell *P*_*k*_, *iMeGroCy* reduces both rates of protein synthesis and time constant of protein degradation during the pre-budded period (*G*_*1*_ phase), with the amount of the reduction increasing according to the parent genealogical age. To this end, both *K*_*2*_ and *τ*_*2*_ are decreased to lower and lower values during G_1_, according to the parent genealogical age. We define 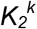 and 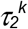 the *K*_*2*_ and *τ*_*2*_ *parameters* for a parent cell with genealogical age ‘*k*’. At the end of timer *T*_*2*_ - coincident with the end of the G_1_ phase and with the onset of the budded phase - the values of 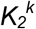 and 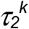 return to the nominal values of *K*_*2*_ and *τ*_*2*_, so that the parent cell *P*_*k*_ grows again with the steady-state exponential rate approximately given by eq.(35). Daughter cells (of any genealogical age) are not affected by such mechanical stress; therefore, their growth parameters are not modified. We model the aforementioned mechanical stress by setting the growth rate λ_*k*_ of a parent cell *P*_*k*_ in G_1_-phase as a percentage of λ, that is the growth rate shared by all daughter cells, as well as all parent cells in the budded phase. In particular, we fixed: λ_*1*_ = 0.85*λ, λ_*2*_ = 0.40*λ, λ_*3*_ = 0.10*λ, λ_*4*_ = 0.05*λ, λ_*5*_ = 0.025*λ, and λ_*k*_ = 0.005* λ for *k* > 5. To obtain the corresponding parameters, *τ*_*2*_ is modified for parent cells *P*_*k*_ within the G_1_-phase, i.e.,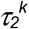 is 50% of *τ*_*2*_.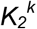 is then computed from 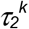 and λ_*k*_, according to eq.(35). Table S3 explicitly reports the values of 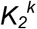 for all the glucose concentrations.

### Simulations of cell populations through *iMeGroCy*

*iMeGroCy* can generate pedigree populations of yeast, distinguishing among the many features of daughter and parent cells.

For any chosen glucose concentration, a preliminary “*long period*” simulation is run, starting from a newborn daughter cell and letting it evolve cycle after cycle, for enough time to reach steady-state conditions (i.e., exponential growth): this way the population properties do not depend on the daughter cell initial conditions. This single germinal line is exploited to build up a population made of 10 germinal lines, each seeded starting from 10 different initial cells (sampled at the end of the “*long period*” simulation) and simulated over time to generate a complete pedigree. The 10 initial cells include daughter or parent cells of different genealogical ages (*k* = 1,2,3,…), different initial protein and ribosome content (P_0_, R_0_) and different positions in the cell cycle at the time of their virtual sampling. We chose 5 daughters (3D_1_, 1D_2_, 1D_3_) and 5 parents (3P_1_, 1P_2_, 1P_3_) accordingly to the daughter/parent cell distributions of the first “*long period*” simulation. In particular, for any chosen glucose concentration, the protein contents of the 10 initial cells, P_0_(D_k_), P_0_(P_k_), *k* = 1,2,3, are sampled from lognormal distributions, with mean values and standard deviations set according to the protein contents distributions of D_k_ and P_k_ cells of the “*long period*” germinal line population in steady-state, grown in the same glucose environment. The selected protein contents of the 10 initial cells are reported in Table S3 for all the glucose concentrations.

Regarding the position in the cell cycle, each germinal line initial cell is supposed to be virtually sampled at Δ minutes from its birth: without loss of generality, we chose Δ = 10*n min, with n = 0, 1, …, 9, thus assuming to have 10 initial cells sampled each 10 min over an average cycle length of 100 min. Such a choice ensures the maximum entropy among the cells and reduces the probability to have unnatural synchronizations. Each pedigree population line is kept proliferating until it reaches steady state, with a final number of N cells each, that is a whole population of 10N cells. Number N is strictly related to the generations required to reach steady-state and may depend on the glucose concentration.

We adopted the following additional rules to simulate the growth of the population:

i. all cells evolve according to the equations described for individual cells;
ii. the lengths of timers *T*_*2*_ and *T*_*B*_ are drawn from a lognormal distribution with average value equal to the values reported in Table S3 and Coefficient of Variation (CV) of 0.05 (equal for daughters and parents of any genealogical age). The CVs (here and in the following items) refer to single-cell variability, providing differences in the properties of individual cells of a given population; the value of the CVs has been set to best fit the protein distributions and is the same for the different periods (to avoid unnecessary complications);
iii. timer *T*_*1b*_ length is drawn from a lognormal distribution with the average value given by a function of the protein content at the onset of *T*_*1b*_ (see eq.(36)) and a given CV of 0.05;
iv. any daughter cell coming at division gives birth to a pair of newborn daughter/parent cells with unitary genealogical age, i.e. *D*_*1*_ and *P*_*1*_; a parent cell *P*_*k*_ gives birth to a pair of newborn daughter/parent cells with incremental genealogical age, i.e. *D*_*k+1*_ and *P*_*k+1*_; see Figure 2C;
v. at cellular division each cell is substituted by a pair of newborn daughter and parent;
vi. the newborn parent cell starts with an initial protein content *P*_*0*_*(P)* drawn from a lognormal distribution with average value given by the critical size *P*_*s*_ achieved by the mother cell at the onset of the budded phase. The CV is taken to be smaller and smaller for newborn parent cells of increasing genealogical age, that is 0.05 for a parent born from a daughter or from a parent *P*_*k*_, k = 1,…,5, while CV = 0.04, 0.03 and 0.02 for a parent born from *P*_*6*_, *P*_*7*_ and *P*_*k*_, k > 7, respectively; the newborn daughter cell starts with an initial protein content *P*_*0*_*(D)* given by the difference of the protein content at division and the newborn parent cell initial size;
vii. the newborn daughter/parent cells start with the initial ribosome content determined (from the overall mother cell ribosome content at division *R*_*CD*_) in order to ensure that the ratios *R*_*0*_*(D)/R*_*CD*_ and *R*_*0*_*(P)/R*_*CD*_ are equal to the ratios *P*_*0*_*(D)/P*_*CD*_ and *P*_*0*_*(P)/P*_*CD*_ respectively.

The outcomes of the population simulations can be clustered into three classes. On the one hand, we have a set of population variables that evolve with time, like (i) the number of cells, (ii) total protein content, (iii) total ribosome content and (iv) the budding index, defined as the fraction of budded cells. According to these four outputs, further outcomes can be derived, such as the population exponential growth (according to which the Mass Duplication Time, MDT, is readily derived as MDT = log(2)/λ) or other time-varying variables, such as the average protein (and ribosome) content. Fig. S1A-D reports some of these evolutions for 2% and 0.05% glucose, with red curves that refer to the average value coming from the 10-line pedigree populations. In Table S4 it can be appreciated the good agreement of the simulated MDT with the experimental values. Other outputs are provided by selecting a set of population cells. To this end, we have considered a time instant when the population under investigation becomes of about 10*N* cells (with *N* dependent on the glucose environment), and computed statistics concerning the whole cycle of these cells, such as the timer lengths, their critical size or their size at birth and at division. All these values are computed in average and standard deviation and can refer to the whole population or to subsets such as daughter or parent cells (or even subsets like *D*_*1*_, *D*_*2*_, *P*_*1*_, *P*_*2*_ cells, etc.). Most of these outputs can be found in Table S4, for different glucose concentrations, and compared to their corresponding experimental values.

The third class of outputs deals with the protein distributions. We have randomly extracted *M* = 50,000 cells from the complete set of 10*N* cells at a given time *t*. The set of *M* cells is “*frozen*” from the population simulation to their actual protein content at the selected time *t*. Each cell is referred to be a daughter or a parent cell, thus distinct daughter and parent protein distributions can be computed. Figure 3B-H reports the protein distributions compared to the corresponding experimental distributions of about 50,000 cells for different glucose and ethanol environments.

### Simulations of cell populations through Hy-*iMeGroCy*

To illustrate our “whole-cell modelling” concept, we used the *iMeGroCy* model as a scaffold for finer molecular plug-ins. To this end, a molecular model of the G_1_/S transition is plugged into the *iMeGroCy*, to replace the hitherto fitted timers describing the G_1_ phase (i.e., T_1_ and T_2_), Figure 6A. The adopted G1_S module ^53^, describes in detail the molecular mechanisms leading to the activation of the G_1_/S regulon. Briefly, the resulting *Hy-iMeGroCy* model works as follows. Synthesis of cyclin Cln3, whose total amount is proportional to the overall protein content (similarly to *GroCy*), links cell growth and molecular events promoting the G_1_/S transition. These events include regulation of Cln3-Cdk1 activity and transcriptional activation of the G_1_/S regulon - driven by SBF and MBF and inhibited by Whi5. The activation of the G_1_/S regulon leads to the synthesis of Cln1, Cln2, Clb5, Clb6, and Nrm1, which play a direct role in later molecular events. The model includes cytoplasm (cyt), endoplasmic reticulum (ER) and nucleus (nuc) sub-cellular compartments. The G_1_/S regulon activation depends on multi-site phosphorylation of Whi5, SBF and MBF catalyzed by Cln-Cdk1 complexes. The T_1_ period lasts from birth up to when 50% of Whi5 has left the nucleus at the end of the multisite phosphorylation period; then T_2_ period starts, and is taken to end when 50% of Sic1 – the inhibitor the Clb-Cdk1 complexes - has left the nucleus following phosphorylation by Cln1,2,3- and Clb5,6-cdk1 complexes.

The G1_S module in ^53^ simulates and predicts single daughter cells or sets of independent daughter cells: to exploit it to make population simulations (of daughter and parent cells, of different genealogical age) is a novelty of the present manuscript, as well as the tuning of its parameters to reproduce different glucose environments.

G1_S simulations in ^53^ refer to cells growing in high glucose. The related parameter values have been here considered as the starting point for the glucose 2% case, with the aim to make *Hy-iMeGroCy* population simulations consistent with *iMeGroCy* population simulations. Only a few of the G1_S module parameters had to be slightly modified from ^53^. The G1_S module parameters tuning procedure has been to best fit timers and protein contents provided by *iMeGroCy* cell chains. The use of small cell chains instead of populations has been considered a good compromise between accuracy and fastness, because a trial and error approach would not be affordable for populations of thousands of cells involving the G1_S module (see ^53^ for the computational burden associated to G1_S module).

The G1_S module parameters that have the same meaning of the corresponding ones in GroCy (for instance, Cln3 and Far1 binding/unbinding coefficients or the initial amount of Far1), are set accordingly to iMeGroCy. Other G1_S model parameters values that do not vary with the genealogical age are taken from ^53^. Table S7 reports, for each glucose concentration, the *Hy-iMeGroCy* parameter values that vary according to the glucose concentration, the type of cell, and the genealogical age. In general, the smallest number of parameters have been changed compared to ^53^, in keeping with the notion that cell cycle parameters are likely to change significantly only at the lowest glucose concentrations. In detail:

- k_14_ (the binding coefficient of the reaction Swi6Swi4 + Whi5 --> Swi6Swi4Whi5) increases, or at least remains constant, when the glucose concentration decreases; it is set to higher values in daughter cells rather than in parent cells; it decreases, or at most it remains constant, by increasing the genealogical age;
- k_21_ (the diffusion coefficient of Whi5 from the nucleus into the cytoplasm) and k_21p_ (the diffusion coefficient of phosphorylated Whi5 from the nucleus into the cytoplasm) decrease, or at most remain constant, when the glucose concentration decreases; they are set to lower values in daughter cells rather than in parent cells; they increase, or at least remain constant, by increasing the genealogical age;
- k_31_ (the binding coefficient of the reaction Clb6 + Sic1 --> Clb6Sic1), k_33_ (the binding coefficient of the reaction Clb6 + Sic1p --> Clb6Sic1p), k_37_ (the binding coefficient of the reaction Clb5 + Sic1 --> Clb5Sic1), k_39_ (the binding coefficient of the reaction Clb5 + Sic1p ->Clb5Sic1p) increase, or at least they remain constant, when the glucose concentration decreases; they are set to higher values in daughter cells rather than in parent cells; they decrease, or at most they remain constant, by increasing the genealogical age;
- k_43_ (the diffusion coefficient of Sic1p out of the nucleus) decreases, or at most, it remains constant, when the glucose concentration decreases; it is set to lower values in daughter cells rather than in parent cells; it increases, or at least it remains constant, by increasing the genealogical age;
- Sic1_tot_ (the initial amount of Sic1) has a lower value for parent cells rather than daughter cells.

Figure 6B-C compares protein distributions for the *Hy-iMeGroCy* to the corresponding experimental ones for different 0.05% and 2% glucose environments. Table S8 reports outputs for the *Hy-iMeGroCy* population simulations for different glucose concentrations.

### Simulations of mutant populations

#### rsa1 - 2% glucose

We increase the MDT by suitably reducing (by about 36%) the protein synthesis rate K_2_ of the WT, CEN.PK - 2% glucose, and we modify the following parameters related to the G_1_ phase varying their values w.r.t. the WT case: Cln3Far1 binding coefficient k_on_ is doubled; parameter W_1_ is slightly increased (by 0.0025%); timer T_2_ length is reduced by 40%. Besides, timer T_B_ length is set equal to the experimental value (i.e. increased by 41% w.r.t. WT). See Table S6 for a summary of input parameters variations. Protein distributions and temporal features are reported in Figure 5A, compared to experimental counterparts.

#### whi5Δ and whi5^4E^ - 2% glucose

For the coarse-grain model, the same input modifications provide a satisfactory fitting for both whi5Δ and whi5^4E^ mutants. Regards to the coarse-grain *iMeGroCy*, we did not vary input growth parameters. Instead, we modified w.r.t. WT the model parameters related to the G_1_ phase as follows: we slightly increased W_1_ of 0.006% and reduced T_2_ of 50%. We also increase timer T_B_ of about 4%. See Table S6 for a summary of input parameters variations. Figure 5B-C reports experimental and simulated protein distributions and temporal features.

The fine-grain model *Hy-iMeGroCy* accounts for the specific *whi5*Δ mutant by simply setting Whi5_tot_ = 0 in the molecular module. For the simulation of the *whi5*^*4E*^ mutant, expressing a protein whose 4 functional sites have been mutated to glutamate that acts as a constitutive phosphomimetic ^53^, we assume that Whi5^4E^ keeps unchanged the possibility to bind to (and therefore inhibit) SBF, but with a lower affinity, that means we reduce the value of k_14_ to 1/10 of the nominal value in Table S7. Moreover, because of the ‘natural’ phosphorylation state of Whi5, soon after the binding of SBF/Whi5, Whi5 is ready to be released without waiting for further phosphorylation(s). However, this happens at a lower rate with respect to the wild type case. See ^53^ for a more complete description. Figure 6D-E reports experimental and simulated protein distributions and temporal features.

#### TM6*

The population growth of mutant TM6* has been simulated under different nutrient conditions, i.e. glucose 2%, 0.1% and ethanol 2% (see Table S5 for input parameters variations). In order to model the TM6* low efficiency of glucose transport we reduced the WT catalytic coefficient of the *MeGro* flux *v*_*hxt*_(*k*_*cat, hxt*_) by a factor of 5. This is consistent with direct in vivo measurements of the glucose uptake capacity for this mutant (Fig. S3 panel I; ^49^). Then, *MeGro* has been run for TM6* mutant providing the following inputs: (i) the experimental glucose concentration 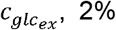 or 0.1%, together with the corresponding experimental value of *F* (setting 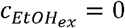); (ii) the experimental concentration 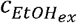 related to ethanol 2% (setting 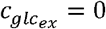), with *F* = 0 and the fixed protein investment *γ*_*hxt*_ = 0.1. The *MeGro* outputs related to the inputs (i) and (ii), reported in Table S5, show the following variations w.r.t. the WT outputs, reported in Table S3: (a) a reduction of both the ribosome-over-protein ratio ρ (∼8% for 0.1% and ∼5% for 2%) and the exponential growth rate λ (∼18% for 0.1% and ∼24% for 2%), as well as a reduction of the protein synthesis rate K_2_ (∼9% for 0.1% and ∼18% for 2%), for case (i); (b) a reduction of both the ribosome-over-protein ratio ρ (∼27%) and the exponential growth rate λ (∼11%), and an increasing of K_2_(∼22%), for case (ii). Finally, the population simulation is performed setting *T*_*B*_ to the experimental value of TM6*. The results on the TM6* simulations are reported in Fig. 4F.

#### hxk2 hxk1

Mutant *hxk2 hxk1* has been simulated under the nutrient conditions 2%, 0.1% glucose and 2% ethanol. To reproduce the *hxk2 hxk1* low efficiency of glycolysis in glucose environment we reduced the WT catalytic coefficient of the flux *v*_*gly*_(*k*_*cat, gly*_) by a factor of 1/2, which is roughly consistent with the reduction in the glucose kinase activity measured in this mutant strain (Fig. S3J). No loss of efficiency is assumed in ethanol 2%. *MeGro* has been run providing the following inputs related to mutant *hxk2 hxk1*: (i) the experimental glucose concentration 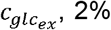 or 0.1%, together with the corresponding experimental value of *F* (setting 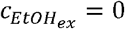); (ii) the experimental concentration 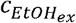 related to ethanol 2% (setting 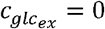), with *F* = 0 and the fixed protein investment *γ*_*hxt*_ =0.01. All other *MeGro* parameters used for the CEN.PK are set according to Table S2. The *hxk2 hxk1* input parameter variations and outputs are reported in Table S5, and show the following variations w.r.t. the WT outputs reported in Table S3: (a) a reduction of both the ribosome-over-protein ratio ρ (∼3% for 0.1% and ∼5% for 2%) and the exponential growth rate λ (∼21% for 0.1% and ∼32% for 2%), as well as a reduction of the protein synthesis rate K_2_ (∼17% for 0.1% and ∼27% for 2%), for input (i); (b) the same pair ρ, λ, as well as the same K_2_, of the WT for the input (ii). Finally, the population simulation is performed setting *T*_*B*_ to the experimental value of *hxk2 hxk1*. The results are reported in Fig. 4G.

### Sensitivity analysis

A sensitivity analysis of GroCy has been performed by changing a single parameter at a time while keeping the remaining model parameters to their nominal value. In particular, each parameter has been moved “forward” and “backward” with respect to its nominal value set for 2% glucose, choosing the number and sizes of the variations based on reasonable “a priori” evaluations or on limits given by the onset of computational problems. For each parameter setting (resulting from each parameter variation) the exponential growth of a yeast population has been simulated (up to ∼50000 cells). From the virtual population we then extracted some statistical indicators related to the protein distribution and to relevant temporal features (as mass duplication time and duration of timers). This analysis links the GroCy model parameters to the selected quantitative features of the cell population.

From the numerical database obtained by the sensitivity analysis, we also deduced a qualitative/quantitative information on the effect produced on each statistical indicator by the “forward” and “backward” variation of the selected model parameter. Let us denote by *p*^*+*^ and *p*^*-*^ the sets of the values chosen for parameter *p*, obtained by varying “forward” and “backward” the parameter *p* with respect to its nominal value *p*_*0*_:

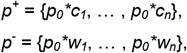

where *c*_*n*_ > … > *c*_*1*_ > 1 and 0 < *w*_*n*_ < … < *w*_*1*_ < 1. To each set *p*^*+*^/*p*^*-*^ we assigned a numerical label *L* identifying the *effect* produced on the *nominal value* (i.e. related to *p*_*0*_) of a given statistical indicator of the virtual population. In particular, we used:

1. *L* = 0, if *all* the values *p*^*+*^/*p*^*-*^ do not substantially change the nominal value of the statistical indicator, i.e. the indicator change belongs to the interval [-5%, 5%];
2. *L* > 0, if *all* the values *p*^*+*^/*p*^*-*^ do not substantially decrease the nominal value of the statistical indicator (below -5%), but *at least one* value of *p*^*+*^/*p*^*-*^ increases the statistical indicator more than 5%;
3. *L* < 0, if *all* the values *p*^*+*^/*p*^*-*^ do not substantially increase the nominal value of the statistical indicator (above 5%), but *at least one* value of *p*^*+*^/*p*^*-*^ decreases the statistical indicator more than -5%.

More in detail, in case 2), if *the strongest* nominal parameter variation in *p*^*+*^/*p*^*-*^ (i.e. *p*_*0*_*\*c*_*n*_*/ p*_*0*_*\*w*_*1*_) increases the nominal statistical indicator more than:

i. 5%, we set *L* = +1;
ii. 25%, we set *L* = +2;
iii. 50%, we set *L* = +3;
iv. 75%, we set *L* = +4;
v. 100%, we set *L* = +5.

Conversely, in case 3), if *the strongest* nominal parameter variation in *p*^*+*^/*p*^*-*^ (i.e. *p*_*0*_*\*c*_*n*_*/ p*_*0*_*\*w*_*1*_) decreases the nominal statistical indicator more than:

i. -5%, we set *L* = -1;
ii. -25%, we set *L* = -2;
iii. -50%, we set *L* = -3;
iv. -75%, we set *L* = -4; (v) -100%, we set *L* = -5.

Figure 2D provides a quick qualitative/quantitative view on how a plus (*p*^*+*^) or minus (*p*^*-*^) alteration of each GroCy input parameter affects the selected statistical indicators. The figure shows the sensitivity heatmap depicting the effects of the up or down parameter variations on the temporal features (mean values of the Mass Duplication Time, <MDT>, and of the G1 periods of daughter, <TG1D>, and parent, <TG1P>, cells) and on the protein distribution features (mean value <P>, standard deviation SD(P) and coefficient of variation CV(P) of the distribution). Supplementary Figure S2 reports an example on how such sensitivity analysis is carried out by varying parameter T_2_.

## SOFTWARE AVAILABILITY

The code to run single cells or populations of *iMeGrocy*, as well as of *Hy-iMeGroCy* can be found at https://drive.google.com/drive/folders/1_JVk42X7Q446YxgAQjjkG4K8B8n7igOa?usp=share_link

## ACKNOWLEDGMENTS

This work was supported by grants to the ISBE-SYSBIO infrastructure, a Ministry of University and Research (MIUR) initiative for the Italian Roadmap of European Strategy Forum on Research Infrastructures (ESFRI) coordinated by L.A. The study also received financial support from the Italian MIUR through grant “Dipartimenti di Eccellenza - 2017” to the University of Milano-Bicocca, Department of Biotechnology and Biosciences and from “ELIXIRxNextGenerationIT” (Code IR0000010)-CUP B53C22001800006. F. P. and S. B. were partially supported by ISBE-SYSIO. The authors are grateful to Prof. E. Boles for providing the *hxt-null* mutant and to late Prof. S. Hohmann for providing the TM6* mutant.

## AUTHOR CONTRIBUTIONS

Conceptualization: L.A., B.T., M.Van.;

Methodology: P.P., F.P., M.Van.;

Software and Visualization: F.P.;

Investigation: S.B., C.A., L.G., P.P., F.P.; I.O., M.Vai

Writing: L.A., M.Van., B.T., P.P., S.B.

---

**Figure 5. A coarse-grained model predicts the protein distributions of mutants in cell growth and cell cycle**

(A-C) Comparison of experimental (red lines) and simulated (blue lines) protein distributions for mutants *rsa1* (A), *whi5*_Δ_ (B) and *whi5*^*4E*^ (C) in glucose 2%. Inserts show temporal parameters for both experimental and simulated populations. All the panels report the experimental distribution of wild-type cells grown in 2% glucose in gray.

(D) Experimental (red bars) and simulated (blue bars) protein synthesis rates for different strains and growth conditions. Data are relative to the wild-type strain cultivated in a 2% glucose medium. Means + SDs from at least two biological replicates are reported.

**Figure 6. A coarse-grained model acts as scaffold for a molecularly detailed module of the G_1_/S transition**

(A) - The G_1_/S transition molecular module plugs in the iMeGroCy model to replace Timers T_1_+T_2_.

(B–E) Protein distributions predicted by the hybrid Hy-iMeGroCy model (blue lines), compared to the related experimental distributions (red lines), for 2% (B) and 0.05% (C) glucose concentrations, and mutants *whi5*_Δ_ (F) and *whi5*^*4E*^ (G). In panels B, D-G, we report the experimental distribution of wild-type cells grown in 2% glucose in gray. Inserts show temporal parameters for both experimental and simulated populations.

