## Supplementary Information for "A modular model integrating metabolism, growth, and cell cycle predicts that fermentation is required to modulate cell size in yeast populations"

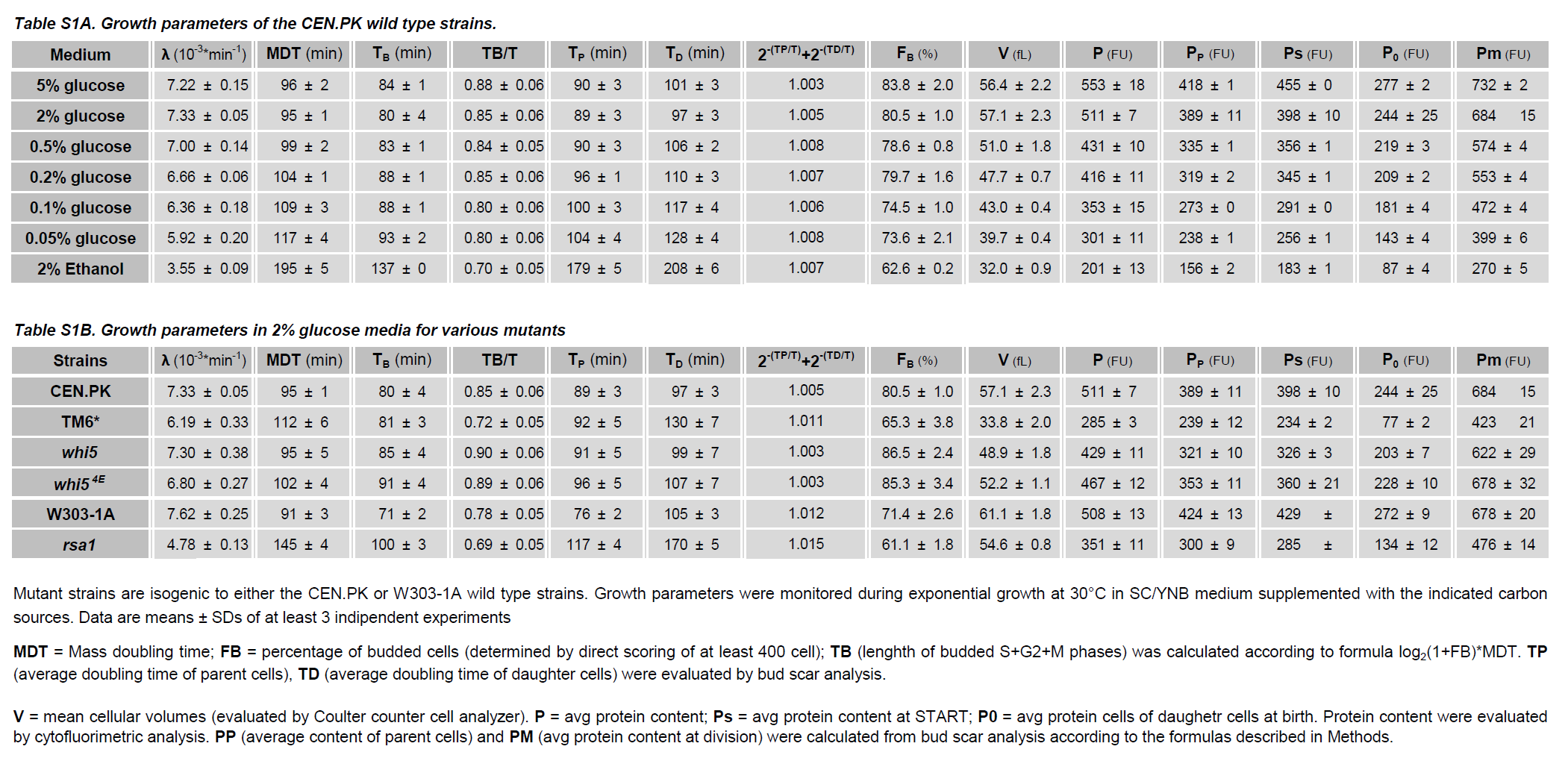

Mutant strains are isogenic to either the CEN.PK or W303-1A wild type strains. Growth parameters were monitored during exponential growth at 30°C in SC/YNB medium supplemented with the indicated carbon sources. Data are means ± SDs of at least 3 independent experiments

**MDT** = Mass doubling time; **FB** = percentage of budded cells (determined by direct scoring of at least 400 cell); **TB** (length of budded S+G2+M phases) was calculated according to formula log_2_(1+FB)*MDT. **TP** (average doubling time of parent cells), **TD** (average doubling time of daughter cells) were evaluated by bud scar analysis.

**V** = mean cellular volumes (evaluated by Coulter counter cell analyzer). **P** = avg protein content; **Ps** = avg protein content at START; **P0** = avg protein cells of daughetr cells at birth. Protein content were evaluated by cytofluorimetric analysis. **P_P_** (average content of parent cells) and **Pm** (avg protein content at division) were calculated from bud scar analysis according to the formulas described in Methods.

**Table S1C. Ethanol yield**

|  | **Ethanol yield** | |
| --- | --- | --- |
| **Medium [glucose]** | ***wild type*** | ***snf3 rgt2 gpa2 gpr1*** |
| **0.05%** | 0.73 (*) | 0.93 |
| **0.1%** | 0.91 * | 0.88 |
| **0.2%** | 1.04 * | 0.81 |
| **0.5%** | 1.07 * | 0.83 |
| **2%** | 1.32 | NA |
| **5%** | 1.50 | 0.78 |

Ethanol yield, i.e., slopes calculated from data in Figure 1(D-E). * indicates statistical significance in a parallelism test (95% confidence) relative to the 5% glucose medium (chosen as the reference growth condition). NA = data not available.

**Table S2. *MeGro* input parameters**

| **Parameter** | **Meas. unit** | **Value** |
| --- | --- | --- |
| ***k_cat,hxt_*** | h^-1^ | 37492 |
| ***k_cat,gly_*** | h^-1^ | 4166 |
| ***k_cat,resp_*** | h^-1^ | 99 |
| ***k_cat,ferm_*** | h^-1^ | 6427 |
| ***k_cat,rib_*** | h^-1^ | 670 |
| ***k_cat,EtOHtr_*** | h^-1^ | 964080 |
| ***k_cat,gly2_*** | h^-1^ | 12497 |
| ***k_cat,resp2_*** | h^-1^ | 221 |
| ***K_M,hxt_*** | mM | 20 |
| ***K_M,gly_*, *K_M,gly2_*** | mM | 0.2 |
| ***K_M,rib_, K_I,pyr_, K_I,glc_*, *K_I2,glc_*** | mM | 1 |
| ***K_M,ADPgly_, K_M,ATPgly2_***  ***K_M,ADPresp_, K_M,ADPresp2,_***  ***K_M,resp_, K_M,ATPrib_*** | mM | 0.5 |
| ***K_M,ferm_*** | mM | 5 |
| ***K_M,EtOHtr_, K_M,resp2_*** | mM | 0.05 |
| ***Nprot/rib*** | #prot/#rib | 79 |
| ***Naa/prot_rib*** | #aa/#prot_rib_ | 157.5 |
| ***Naa/prot*** | #aa/#prot | 441.5 |

**Table S3 – *iMeGroCy* input parameters**

| **GroCy inputs (computed from MeGro)** | | | | | | | | |
| --- | --- | --- | --- | --- | --- | --- | --- | --- |
| **Parameters** | **Meas. unit** | **EtOH 2%** | **glc 0.05%** | **glc 0.1%** | **glc 0.2%** | **glc 0.5%** | **glc 2%** | **glc 5%** |
|  |  | $\boldsymbol{F}\boldsymbol{=0}$ | $\boldsymbol{F}\boldsymbol{=0.804}$ | $\boldsymbol{F}\boldsymbol{=}\mathbf{0.883}$ | $\boldsymbol{F}\boldsymbol{=0.928}$ | $\boldsymbol{F}\boldsymbol{=0.958}$ | $\boldsymbol{F}\boldsymbol{=}\mathbf{0.974}$ | $\boldsymbol{F}\boldsymbol{=0.977}$ |
|  |  | $\boldsymbol{\gamma}_{\boldsymbol{hxt}}\boldsymbol{=0.01}$ | **free invest.** | **free invest.** | **free invest.** | **free invest.** | **free invest.** | **free invest.** |
| ***ρ*** | rib/aa | 8.327e-06 | 1.964e-05 | 2.015e-05 | 2.050e-05 | 2.075e-05 | 2.084e-05 | 2.085e-05 |
| ***K****_2_* | aa/rib/min | 526.80 | 304.17 | 317.17 | 332.37 | 351.10 | 367.69 | 372.07 |
| ***K****_2_^1^* | aa/rib/min | 493.82 | 278.06 | 288.62 | 301.21 | 316.91 | 330.93 | 334.65 |
| ***K****_2_^2^* | aa/rib/min | 274.77 | 148.82 | 153.33 | 158.96 | 166.15 | 172.67 | 174.41 |
| ***K****_2_^3^* | aa/rib/min | 128.74 | 62.66 | 63.14 | 64.12 | 65.64 | 67.16 | 67.58 |
| ***K****_2_^4^* | aa/rib/min | 104.40 | 48.30 | 48.11 | 48.32 | 48.89 | 49.58 | 49.78 |
| ***K****_2_^5^* | aa/rib/min | 92.23 | 41.12 | 40.60 | 40.42 | 40.51 | 40.78 | 40.87 |
| ***K****_2_^s^, s > 5* | aa/rib/min | 82.49 | 35.37 | 34.58 | 34.09 | 33.81 | 33.75 | 33.75 |
| **GroCy inputs (tuned by the modeler)** | | | | | | | | |
| **Parameters** | **Meas. unit** | **EtOH 2%** | **glc 0.05%** | **glc 0.1%** | **glc 0.2%** | **glc 0.5%** | **glc 2%** | **glc 5%** |
| ***Far1(0)*** | molec | 110 | 170 | 240 | 240 | 240 | 240 | 240 |
| ***Cln3****_nuc_****(0)*** | molec | 0 | 0 | 0 | 0 | 0 | 0 | 0 |
| ***Cln3Far1(0)*** | molec | 0 | 0 | 0 | 0 | 0 | 0 | 0 |
| ***Far1****_reset_* | molec | 110 | 170 | 240 | 240 | 240 | 240 | 240 |
| ***K****_1_* | min^-1^ | 1 | 1 | 1 | 1 | 1 | 1 | 1 |
| ***τ****_1_* | min | 4000 | 4000 | 4000 | 4000 | 4000 | 4000 | 4000 |
| ***τ****_2_* | min | 3000 | 3000 | 3000 | 3000 | 3000 | 3000 | 3000 |
| ***τ****_2_^s^, s ≥ 1* | min | 1500 | 1500 | 1500 | 1500 | 1500 | 1500 | 1500 |
| ***k****_on_* | (molec/L)^-1^/min | 1.63e-15 | 1.63e-15 | 1.63e-15 | 1.63e-15 | 1.63e-15 | 1.63e-15 | 1.63e-15 |
| ***k****_off_* | min^-1^ | 25 | 25 | 25 | 25 | 25 | 25 | 25 |
| ***h*** | - | 0.07 | 0.07 | 0.07 | 0.07 | 0.07 | 0.07 | 0.07 |
| ***H*** | aa/L | 6.18e23 | 7.09e23 | 7.09e23 | 7.09e23 | 7.09e23 | 7.09e23 | 7.09e23 |
| ***n****_F_* | - | 10 | 10 | 10 | 10 | 10 | 10 | 10 |
| $\bar{\eta}$ | min^-1^ | 1 | 1 | 1 | 1 | 1 | 1 | 1 |
| ***Θ*** | molec/aa | 2.66e-8 | 3.02e-8 | 3.02e-8 | 3.02e-8 | 3.02e-8 | 3.02e-8 | 3.02e-8 |
| ***k****_cn_* | min^-1^ | 1.5 | 1.5 | 1.5 | 1.5 | 1.5 | 1.5 | 1.5 |
| ***k****_nc_* | min^-1^ | 0.6 | 0.6 | 0.6 | 0.6 | 0.6 | 0.6 | 0.6 |
| ***T****_1b,min_* | min | 15 | 1 | 1 | 1 | 1 | 1 | 1 |
| ***W****_0_* | min | 1466 | 1496 | 1503 | 1503 | 1503 | 1503 | 1503 |
| ***W****_1_* | min | 62.1 | 62.1 | 62.1 | 62.1 | 62.1 | 62.1 | 62.1 |
| ***T****_2_* | min | 35 | 10 | 10 | 10 | 10 | 10 | 10 |
| ***T****_B_ (simulating strain W303 in Fig.3)* | min | - | 90 | 90 | 90 | 85 | 85 | 85 |
| ***T****_B_ (simulating CEN.PK mutants in Fig.4)* | min | 133 | - | 87 | - | - | 81 | - |
| **Protein content for the initial cells of the 10 germinal lines providing the *iMeGroCy* population**  **(tuned by the modeler)** | | | | | | | | |
| ***P_0_(D_1_)_gl1_*** | aa | 1.46e10 | 1.46e10 | 2.10e10 | 1.99e10 | 2.54e10 | 3.03e10 | 3.76e10 |
| ***P_0_(D_1_)_gl2_*** | aa | 1.15e10 | 1.15e10 | 2.20e10 | 2.30e10 | 2.21e10 | 3.18e10 | 2.71e10 |
| ***P_0_(D_1_)_gl3_*** | aa | 1.03e10 | 1.03e10 | 1.86e10 | 2.73e10 | 2.44e10 | 2.68e10 | 3.37e10 |
| ***P_0_(D_2_)_gl4_*** | aa | 1.48e10 | 1.48e10 | 2.67e10 | 2.85e10 | 3.01e10 | 3.14e10 | 2.77e10 |
| ***P_0_(D_3_)_gl5_*** | aa | 1.66e10 | 1.66e10 | 2.40e10 | 2.64e10 | 2.80e10 | 3.13e10 | 3.50e10 |
| ***P_0_(P_1_)_gl6_*** | aa | 1.62e10 | 1.62e10 | 2.18e10 | 2.73e10 | 3.04e10 | 3.65e10 | 4.03e10 |
| ***P_0_(P_1_)_gl7_*** | aa | 2.08e10 | 2.08e10 | 2.38e10 | 3.06e10 | 3.26e10 | 3.57e10 | 3.51e10 |
| ***P_0_(P_1_)_gl8_*** | aa | 1.92e10 | 1.92e10 | 2.29e10 | 2.89e10 | 3.39e10 | 3.32e10 | 3.73e10 |
| ***P_0_(P_2_)_gl9_*** | aa | 2.63e10 | 2.63e10 | 2.77e10 | 2.95e10 | 3.13e10 | 3.87e10 | 3.35e10 |
| ***P_0_(P_3_)_gl10_*** | aa | 2.49e10 | 2.49e10 | 3.00e10 | 3.52e10 | 3.11e10 | 3.95e10 | 4.42e10 |

**Table S4 - *iMeGroCy* population output parameters. Light cyan rows refer to control experimental data**

| **Parameters** | **Meas. Unit** | **EtOH 2%** | **glc 0.05%** | **glc 0.1%** | **glc 0.2%** | **glc 0.5%** | **glc 2%** | **glc 5%** |
| --- | --- | --- | --- | --- | --- | --- | --- | --- |
| ***MDT*** | min | 187 | 131 | 121 | 112 | 104 | 99 | 98 |
| ***MDT*** | min | 195 ± 5 | 117 ±4 | 109 ± 3 | 104 ±1 | 99 ± 2 | 95 ± 1 | 96 ± 2 |
| **2^-^*^TD/MDT^* + 2^-^*^TP/MDT^*** | - | 1.0000 | 0.996 | 0.999 | 1.000 | 0.999 | 0.999 | 0.997 |
| **2^-^*^TD/MDT^* + 2^-^*^TP/MDT^*** | - | 1.007 ± 0.003 | 1.008 ± 0.004 | 1.006 ± 0.001 | 1.007 ± 0.002 | 1.008 ± 0.002 | 1.003 ± 0.002 | 1.003 ± 0.000 |
| ***T_G1_(D)*** | min | 58 | 55.6 ± 12.6 | 41.8 ± 10.5 | 28.8 ± 8.5 | 25.2 ± 8.0 | 15.9 ± 5.8 | 14.1 ± 4.7 |
| ***T_G1_(D)*** | min | 71 | 35 | 29 | 22 | 23 | 21 | 17 |
| ***T_G1_(P)*** | min | 52 | 28.5 ± 9.5 | 20.7 ± 7.2 | 14.9 ± 4.6 | 13.3 ± 3.4 | 11.5 ± 1.3 | 11.3 ± 0.9 |
| ***T_G1_(P)*** | min | 42 | 11 | 12 | 8 | 7 | 9 | 6 |
| ***T_B_*** | min | 131.80 | 90 ± 5 | 90 ± 5 | 90 ± 5 | 85 ± 4 | 85 ± 4 | 85 ± 4 |
| ***T_B_*** | min | 137 ± 1 | 93 ± 2 | 88 ± 1 | 88 ± 1 | 83 ± 1 | 82 ± 2 | 84 ± 1 |
| ***T_D_*** | min | 190 | 146 ± 13 | 132 ± 12 | 119 ± 10 | 110 ± 9 | 101 ± 7 | 99 ± 6 |
| ***T_D_*** | min | 208 ± 6 | 128 ± 4 | 117 ± 4 | 110 ± 3 | 106 ± 2 | 103 ± 1 | 101 ± 1 |
| ***T_P_*** | min | 183 | 119 ± 11 | 111 ± 9 | 105 ± 6 | 99 ± 5 | 97 ± 4 | 97 ± 4 |
| ***T_P_*** | min | 179 ± 5 | 104 ± 4 | 100 ± 3 | 96 ± 1 | 90 ± 3 | 91 ± 1 | 90 ± 3 |
| ***<P>/<P>_2%_*** | - | 0.51 | 0.53 | 0.71 | 0.83 | 0.87 | 1.0 | 1.08 |
| ***<P>/<P>_2%_*** | - |  | 0.60 | 0.70 | 0.79 | 0.86 | 1.0 | 1.13 |

**Table S5 – *iMeGroCy* input/output parameter variations for mutants TM6* and *hxk2 hxk1***

| **Parameters** | **TM6*** | | | **hxk2 hxk1** | | |
| --- | --- | --- | --- | --- | --- | --- |
|  | **EtOH 2%** | **glc 0.1%** | **glc 2%** | **EtOH 2%** | **glc 0.1%** | **glc 2%** |
|  | $\boldsymbol{F}\boldsymbol{=0}$ | $\boldsymbol{F}\boldsymbol{=}\mathbf{0.460}$ | $\boldsymbol{F}\boldsymbol{=}\mathbf{0.608}$ | $\boldsymbol{F}\boldsymbol{=0}$ | $\boldsymbol{F}\boldsymbol{=}\mathbf{0.242}$ | $\boldsymbol{F}\boldsymbol{=}\mathbf{0.511}$ |
|  | $\boldsymbol{\gamma}_{\boldsymbol{hxt}}\boldsymbol{=0.1}$ | **free invest.** | **free invest.** | $\boldsymbol{\gamma}_{\boldsymbol{hxt}}\boldsymbol{=0.01}$ | **free invest.** | **free invest.** |
|  | **k_cat,hxt_= k_cat,hxt_^nom^/5** | **k_cat,hxt_= k_cat,hxt_^nom^/5** | **k_cat,hxt_= k_cat,hxt_^nom^/5** | **No MeGro parameter variations** | **k_cat,gly_= k_cat,gly_^nom^/2,**  **k_cat,gly2_= k_cat,gly2_^nom^/2** | **k_cat,gly_= k_cat,gly_^nom^/2,**  **k_cat,gly2_= k_cat,gly2_^nom^/2** |
| ***ρ*** | 6.085e-06 | 1.850e-05 | 1.980e-05 | 8.327e-06 | 1.945e-05 | 1.986e-05 |
| ***K****_2_* | 645.14 | 287.51 | 300.10 | 526.80 | 263.66 | 267.76 |
| ***K****_2_^1^* | 611.37 | 265.11 | 274.45 | 493.82 | 243.82 | 246.90 |
| ***K****_2_^2^* | 345.70 | 143.84 | 146.98 | 274.77 | 132.89 | 133.96 |
| ***K****_2_^3^* | 168.59 | 62.99 | 62.00 | 128.74 | 58.93 | 58.67 |
| ***K****_2_^4^* | 139.08 | 49.52 | 47.84 | 104.40 | 46.60 | 46.12 |
| ***K****_2_^5^* | 124.32 | 42.78 | 40.76 | 92.23 | 40.44 | 39.85 |
| ***K****_2_^s^, s > 5* | 112.51 | 37.39 | 35.09 | 82.49 | 35.51 | 34.83 |
| ***T****_B_* | 146 | 87 | 79 | 137 | 103 | 100 |

**Table S6 – iMeGroCy input/output parameter variations for mutants *rsa1*, *whi5* and *whi5^4E^***

| **Parameters** | **rsa1** | **whi5^4E^/whi5** |
| --- | --- | --- |
|  | **glc 2%** | **glc 2%** |
| ***K****_2_* | 255.19 | 400.98 |
| ***K****_2_^1^* | 236.04 | 359.96 |
| ***K****_2_^2^* | 128.69 | 187.01 |
| ***K****_2_^3^* | 57.13 | 71.70 |
| ***K****_2_^4^* | 45.20 | 52.49 |
| ***K****_2_^5^* | 39.23 | 42.88 |
| ***K****_2_^s^, s > 5* | 34.46 | 35.19 |
| ***T****_B_* | 100 | 74 |
| ***T****_2_* | 4 | 5 |
| ***W****_1_* | 62.3 | 62.5 |
| ***k****_on_* | 3.26e-15 | - |

**Table S7 –** ***G1_S* module input parameters**

| **Parameters** | **Meas. unit** | **glc 0.05%** | **glc 0.2%** | **glc 2%** | **glc 5%** |
| --- | --- | --- | --- | --- | --- |
| ***k_14_(D_1_)*** | (molec/L)^-1^min^-1^ | 5.60e-14 | 4.48e-14 | 1.60e-14 | 1.57e-14 |
| ***k_14_(D_2_)*** | (molec/L)^-1^min^-1^ | 5.28e-14 | 4.32e-14 | 7.12e-15 | 5.28e-15 |
| ***k_14_(D_3_)*** | (molec/L)^-1^min^-1^ | 5.12e-14 | 2.72e-14 | 3.14e-15 | 2.32e-15 |
| ***k_14_(D_4_)*** | (molec/L)^-1^min^-1^ | 5.12e-14 | 2.4e-14 | 3.14e-15 | 2.03e-15 |
| ***k_14_(D_5_)*** | (molec/L)^-1^min^-1^ | 4.96e-14 | 2.4e-14 | 2.85e-15 | 1.92e-15 |
| ***k_14_(D_6_)*** | (molec/L)^-1^min^-1^ | 4.96e-14 | 2.4e-14 | 2.70e-15 | 1.84e-15 |
| ***k_14_(D_k_), k > 6*** | (molec/L)^-1^min^-1^ | 4.96e-14 | 2.4e-14 | 2.67e-15 | 1.79e-15 |
| ***k_14_(P_k_), k ≥ 1*** | (molec/L)^-1^min^-1^ | 3.20e-14 | 1.68e-15 | 1.68e-15 | 1.68e-15 |
| ***k_21_(D_1_)*** | min^-1^ | 0.0024 | 0.0062 | 0.11 | 0.31 |
| ***k_21_(D_2_)*** | min^-1^ | 0.0030 | 0.0105 | 2.80 | 4.76 |
| ***k_21_(D_3_)*** | min^-1^ | 0.0033 | 0.0136 | 3.50 | 4.76 |
| ***k_21_(D_4_)*** | min^-1^ | 0.0034 | 0.0145 | 4.20 | 5.18 |
| ***k_21_(D_5_)*** | min^-1^ | 0.0035 | 0.0151 | 4.20 | 5.60 |
| ***k_21_(D_6_)*** | min^-1^ | 0.0035 | 0.0153 | 4.20 | 5.74 |
| ***k_21_(D_k_), k > 6*** | min^-1^ | 0.0035 | 0.0153 | 4.20 | 5.74 |
| ***k_21_(P_1_)*** | min^-1^ | 0.0046 | 0.0336 | 6.02 | 6.02 |
| ***k_21_(P_2_)*** | min^-1^ | 0.0100 | 2.03 | 6.02 | 6.02 |
| ***k_21_(P_3_)*** | min^-1^ | 0.0161 | 4.48 | 6.02 | 6.02 |
| ***k_21_(P_4_)*** | min^-1^ | 0.0196 | 5.53 | 6.02 | 6.02 |
| ***k_21_(P_5_)*** | min^-1^ | 0.0210 | 6.02 | 6.02 | 6.02 |
| ***k_21_(P_6_)*** | min^-1^ | 0.0210 | 6.02 | 6.02 | 6.02 |
| ***k_21_(P_k_), k > 6*** | min^-1^ | 0.0210 | 6.02 | 6.02 | 6.02 |
| ***k_21p_(D_1_)*** | min^-1^ | 0.024 | 0.062 | 1.1 | 3.08 |
| ***k_21p_(D_2_)*** | min^-1^ | 0.030 | 0.105 | 28 | 47.6 |
| ***k_21p_(D_3_)*** | min^-1^ | 0.033 | 0.136 | 35 | 47.6 |
| ***k_21p_(D_4_)*** | min^-1^ | 0.034 | 0.145 | 42 | 51.8 |
| ***k_21p_(D_5_)*** | min^-1^ | 0.035 | 0.151 | 42 | 56.0 |
| ***k_21p_(D_6_)*** | min^-1^ | 0.035 | 0.153 | 42 | 57.4 |
| ***k_21p_(D_k_), k > 6*** | min^-1^ | 0.035 | 0.153 | 42 | 57.4 |
| ***k_21p_(P_1_)*** | min^-1^ | 0.046 | 0.336 | 60.2 | 60.2 |
| ***k_21p_(P_2_)*** | min^-1^ | 0.100 | 20.3 | 60.2 | 60.2 |
| ***k_21p_(P_3_)*** | min^-1^ | 0.161 | 44.8 | 60.2 | 60.2 |
| ***k_21p_(P_4_)*** | min^-1^ | 0.196 | 55.3 | 60.2 | 60.2 |
| ***k_21p_(P_5_)*** | min^-1^ | 0.210 | 60.2 | 60.2 | 60.2 |
| ***k_21p_(P_6_)*** | min^-1^ | 0.210 | 60.2 | 60.2 | 60.2 |
| ***k_21p_(P_k_), k > 6*** | min^-1^ | 0.210 | 60.2 | 60.2 | 6.02 |
| ***k_31_(D_1_)*** | min^-1^ | 2.88e-15 | 2.18e-15 | 1e-15 | 1e-15 |
| ***k_31_(D_2_)*** | min^-1^ | 2.3e-15 | 1.85e-15 | 1e-15 | 8e-16 |
| ***k_31_(D_3_)*** | min^-1^ | 2.23e-15 | 1.8e-15 | 8e-16 | 2e-16 |
| ***k_31_(D_4_)*** | min^-1^ | 2.21e-15 | 1.6e-15 | 3.5e-16 | 2e-16 |
| ***k_31_(D_5_)*** | min^-1^ | 2.21e-15 | 1.59e-15 | 2.5e-16 | 2e-16 |
| ***k_31_(D_6_)*** | min^-1^ | 2.1e-15 | 1.59e-15 | 2.09e-16 | 2e-16 |
| ***k_31_(D_k_), k > 6*** | min^-1^ | 2.08e-15 | 1.58e-15 | 2e-16 | 2e-16 |
| ***k_31_(P_1_)*** | min^-1^ | 2e-15 | 1.56e-15 | 2e-16 | 2e-16 |
| ***k_31_(P_2_)*** | min^-1^ | 2e-15 | 1.56e-15 | 2e-16 | 2e-16 |
| ***k_31_(P_k_), k > 2*** | min^-1^ | 2e-15 | 2e-16 | 2e-16 | 2e-16 |
| ***k_33_(D_1_)*** | min^-1^ | 2.88e-16 | 2.18e-16 | 1e-16 | 1e-16 |
| ***k_33_(D_2_)*** | min^-1^ | 2.3e-16 | 1.85e-16 | 1e-16 | 8e-17 |
| ***k_33_(D_3_)*** | min^-1^ | 2.23e-16 | 1.8e-16 | 8e-17 | 2e-17 |
| ***k_33_(D_4_)*** | min^-1^ | 2.21e-16 | 1.6e-16 | 3.5e-17 | 2e-17 |
| ***k_33_(D_5_)*** | min^-1^ | 2.21e-16 | 1.59e-16 | 2.5e-17 | 2e-17 |
| ***k_33_(D_6_)*** | min^-1^ | 2.1e-16 | 1.59e-16 | 2.09e-17 | 2e-17 |
| ***k_33_(D_k_), k > 6*** | min^-1^ | 2.08e-16 | 1.58e-16 | 2e-17 | 2e-17 |
| ***k_33_(P_1_)*** | min^-1^ | 2e-16 | 1.56e-16 | 2e-17 | 2e-17 |
| ***k_33_(P_2_)*** | min^-1^ | 2e-16 | 1.56e-16 | 2e-17 | 2e-17 |
| ***k_33_(P_k_), k > 2*** | min^-1^ | 2e-16 | 2e-17 | 2e-17 | 2e-17 |
| ***k_37_(D_1_)*** | min^-1^ | 2.88e-15 | 2.18e-15 | 1e-15 | 1e-15 |
| ***k_37_(D_2_)*** | min^-1^ | 2.3e-15 | 1.85e-15 | 1e-15 | 8e-16 |
| ***k_37_(D_3_)*** | min^-1^ | 2.23e-15 | 1.8e-15 | 8e-16 | 2e-16 |
| ***k_37_(D_4_)*** | min^-1^ | 2.21e-15 | 1.6e-15 | 3.5e-16 | 2e-16 |
| ***k_37_(D_5_)*** | min^-1^ | 2.21e-15 | 1.59e-15 | 2.5e-16 | 2e-16 |
| ***k_37_(D_6_)*** | min^-1^ | 2.1e-15 | 1.59e-15 | 2.09e-16 | 2e-16 |
| ***k_37_(D_k_), k > 6*** | min^-1^ | 2.08e-15 | 1.58e-15 | 2e-16 | 2e-16 |
| ***k_37_(P_1_)*** | min^-1^ | 2e-15 | 1.56e-15 | 2e-16 | 2e-16 |
| ***k_37_(P_2_)*** | min^-1^ | 2e-15 | 1.56e-15 | 2e-16 | 2e-16 |
| ***k_37_(P_k_), k > 2*** | min^-1^ | 2e-15 | 2e-16 | 2e-16 | 2e-16 |
| ***k_39_(D_1_)*** | min^-1^ | 2.88e-16 | 2.18e-16 | 1e-16 | 1e-16 |
| ***k_39_(D_2_)*** | min^-1^ | 2.3e-16 | 1.85e-16 | 1e-16 | 8e-17 |
| ***k_39_(D_3_)*** | min^-1^ | 2.23e-16 | 1.8e-16 | 8e-17 | 2e-17 |
| ***k_39_(D_4_)*** | min^-1^ | 2.21e-16 | 1.6e-16 | 3.5e-17 | 2e-17 |
| ***k_39_(D_5_)*** | min^-1^ | 2.21e-16 | 1.59e-16 | 2.5e-17 | 2e-17 |
| ***k_39_(D_6_)*** | min^-1^ | 2.1e-16 | 1.59e-16 | 2.09e-17 | 2e-17 |
| ***k_39_(D_k_), k > 6*** | min^-1^ | 2.08e-16 | 1.58e-16 | 2e-17 | 2e-17 |
| ***k_39_(P_1_)*** | min^-1^ | 2e-16 | 1.56e-16 | 2e-17 | 2e-17 |
| ***k_39_(P_2_)*** | min^-1^ | 2e-16 | 1.56e-16 | 2e-17 | 2e-17 |
| ***k_39_(P_k_), k > 2*** | min^-1^ | 2e-16 | 2e-17 | 2e-17 | 2e-17 |
| ***k_43_(D_1_)*** | min^-1^ | 0.0406 | 0.0692 | 0.308 | 0.413 |
| ***k_43_(D_2_)*** | min^-1^ | 0.0435 | 0.1 | 1.12 | 4.9 |
| ***k_43_(D_3_)*** | min^-1^ | 0.0462 | 0.1194 | 31.5 | 56 |
| ***k_43_(D_4_)*** | min^-1^ | 0.0473 | 0.1215 | 42 | 56 |
| ***k_43_(D_5_)*** | min^-1^ | 0.0476 | 0.1246 | 49 | 56 |
| ***k_43_(D_6_)*** | min^-1^ | 0.0476 | 0.1267 | 54.6 | 56 |
| ***k_43_(D_k_), k > 6*** | min^-1^ | 0.0476 | 0.1267 | 56 | 56 |
| ***k_43_(P_1_)*** | min^-1^ | 0.0512 | 0.1526 | 56 | 56 |
| ***k_43_(P_2_)*** | min^-1^ | 0.0945 | 1.12 | 56 | 56 |
| ***k_43_(P_3_)*** | min^-1^ | 0.1344 | 38.5 | 56 | 56 |
| ***k_43_(P_4_)*** | min^-1^ | 0.1540 | 56 | 56 | 56 |
| ***k_43_(P_5_)*** | min^-1^ | 0.1603 | 56 | 56 | 56 |
| ***k_43_(P_6_)*** | min^-1^ | 0.1603 | 56 | 56 | 56 |
| ***k_43_(P_k_), k > 6*** | min^-1^ | 0.1603 | 56 | 56 | 56 |
| ***Sic1_tot_(D_k_), k≥1*** | min^-1^ | 190 | 190 | 190 | 190 |
| ***Sic1_tot_(P_k_), k≥1*** | min^-1^ | 140 | 140 | 140 | 140 |

**TABLE S8 - *HyG1_S-iMeGroCy* population output parameters. Light cyan rows refer to control experimental data**

| **Parameters** | **Meas. Unit** | **glc 0.05%** | **glc 0.2%** | **glc 2%** | **glc 5%** |
| --- | --- | --- | --- | --- | --- |
| ***MDT*** | Min | 130.8 | 110.9 | 97.4 | 96.3 |
| ***MDT*** | Min | 117 | 104 | 97 | 96 |
| **2^-^*^TD/MDT^* +**  **2^-^*^TP/MDT^*** | - | 0.989 | 0.990 | 0.989 | 0.986 |
| **2^-^*^TD/MDT^* +**  **2^-^*^TP/MDT^*** | - | 1.008 | 1.007 | 1.003 | 1.003 |
| ***T_G1_(D)*** | min | 54.5 ± 8.9 | 29.2 ± 4.7 | 16.6 ± 3.0 | 15.3 ± 2.8 |
| ***T_G1_(D)*** | min | 35 | 22 | 21 | 17 |
| ***T_G1_(P)*** | min | 31.8 ± 7.6 | 16.0 ± 4.1 | 11.2 ± 0.7 | 11.2 ± 0.7 |
| ***T_G1_(P)*** | min | 11 | 8 | 9 | 6 |
| ***T_B_*** | min | 90.0 ± 2.7 | 90.1 ± 2.7 | 85.1 ± 2.6 | 85.1 ± 2.6 |
| ***T_B_*** | min | 93 ± 2 | 88 ± 1 | 82 ± 2 | 84 ± 1 |
| ***T_D_*** | min | 144.5 ± 9.3 | 119.2 ± 5.4 | 101.7 ± 4.0 | 100.4 ± 3.8 |
| ***T_D_*** | min | 128 ± 1 | 110 ± 0 | 103 ± 1 | 101 ± 0 |
| ***T_P_*** | min | 121.8 ± 8.1 | 106.1 ± 4.9 | 96.3 ± 2.7 | 96.3 ± 2.6 |
| ***T_P_*** | min | 104 ± 0 | 96 ± 0 | 91 ± 1 | 90 ± 0 |
| ***F_G1*_*** | % | 10.6 ± 0.9 | 9.8 ± 0.4 | 9.7 ± 0.7 | 9.6 ± 0.4 |
| ***<P>/<P>_2%_*** | - | 0.55 | 0.92 | 1.0 | 1.11 |
| ***<P>/<P>_2%_*** | - | 0.60 | 0.79 | 1.0 | 1.13 |

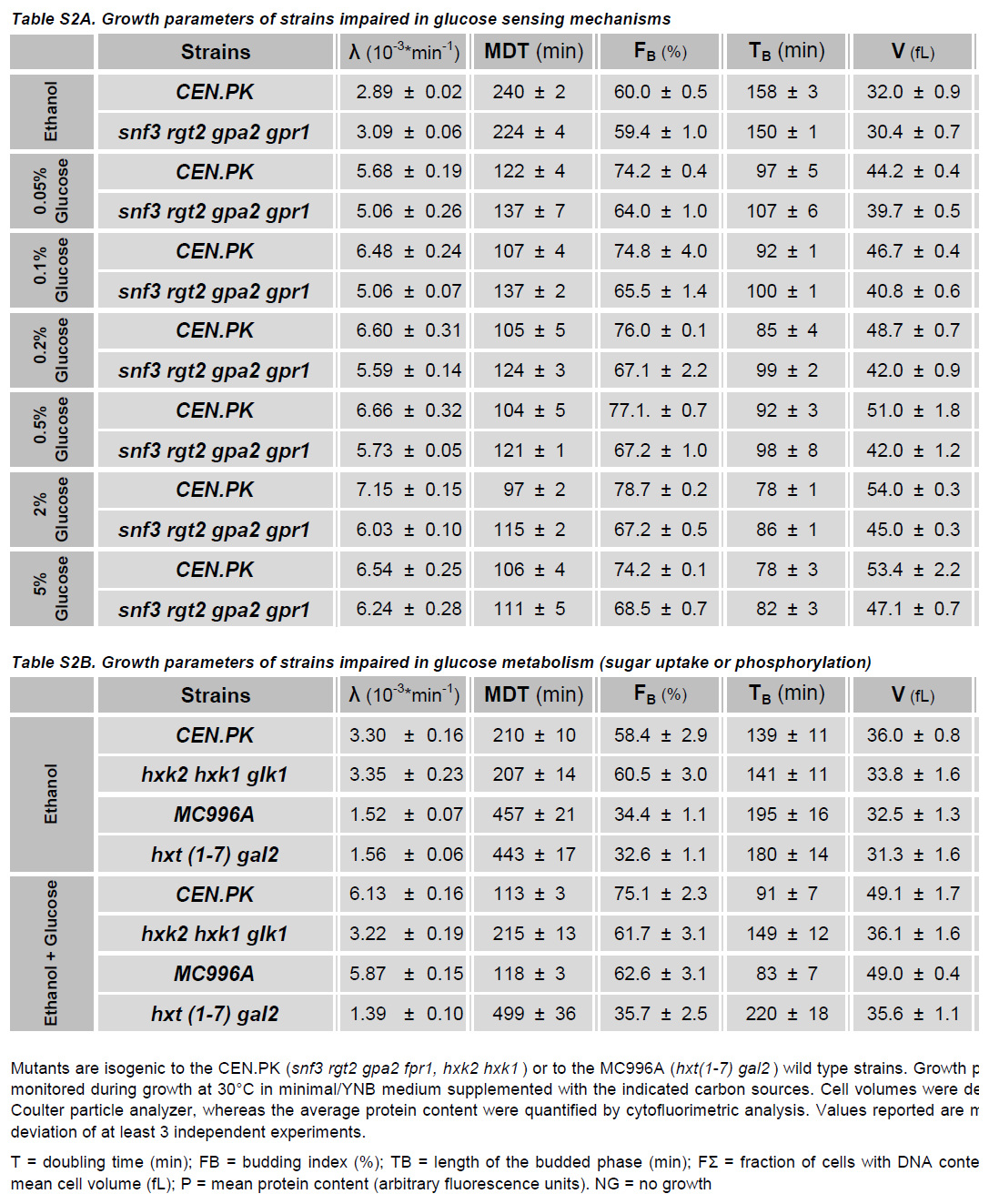
**Table S9. Growth parameters of strains impaired in glucose sensing mechanisms**

Mutant strains are isogenic to either the CEN.PK wild type strains. Growth parameters were monitored during exponential growth at 30°C in SC/YNB medium supplemented with the indicated carbon sources. Data are means ± SDs of at least 3 independent experiments.

**λ** = growth rate; **MDT** = Mass doubling time; **FB** = percentage of budded cells (determined by direct scoring of at least 400 cell); **TB** (length of budded S+G2+M phases) was calculated according to formula

log_2_(1+FB)*MDT.

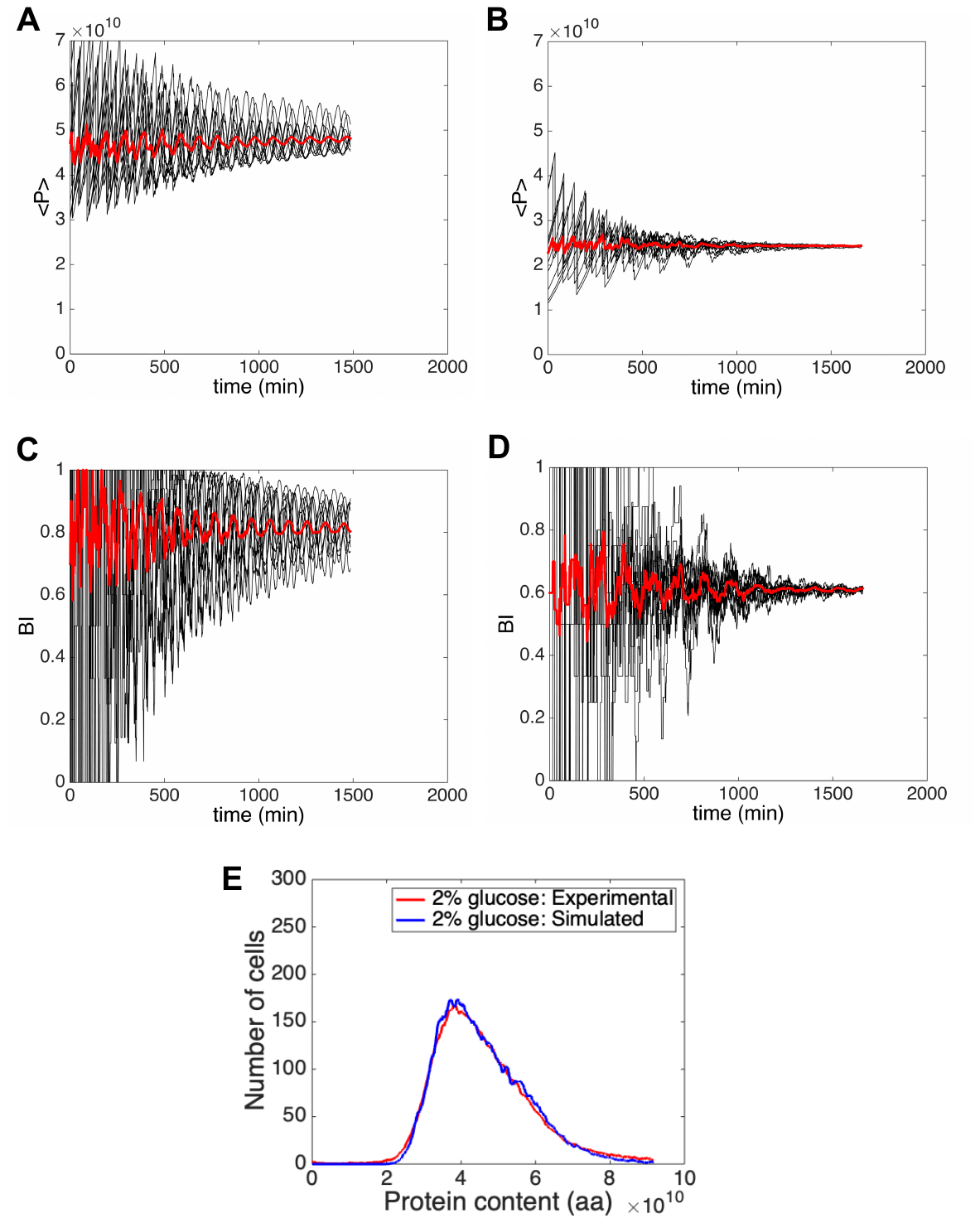
**Supplementary Figure S1 – Simulated parameters in evolving and steady state populations** **- A, B:** time course of the average protein content, for fast (glucose 2%, panel A) and slow (glucose 0.05%, panel B) growth conditions. **C, D:** time course of the budding index, for fast (glucose 2%, panel C) and slow (glucose 0.05%, panel D) growth conditions. The convergence rate depends on the kind of population meta-parameters. **E:** simulated vs experimental protein distribution of a yeast population growing in 2% glucose.

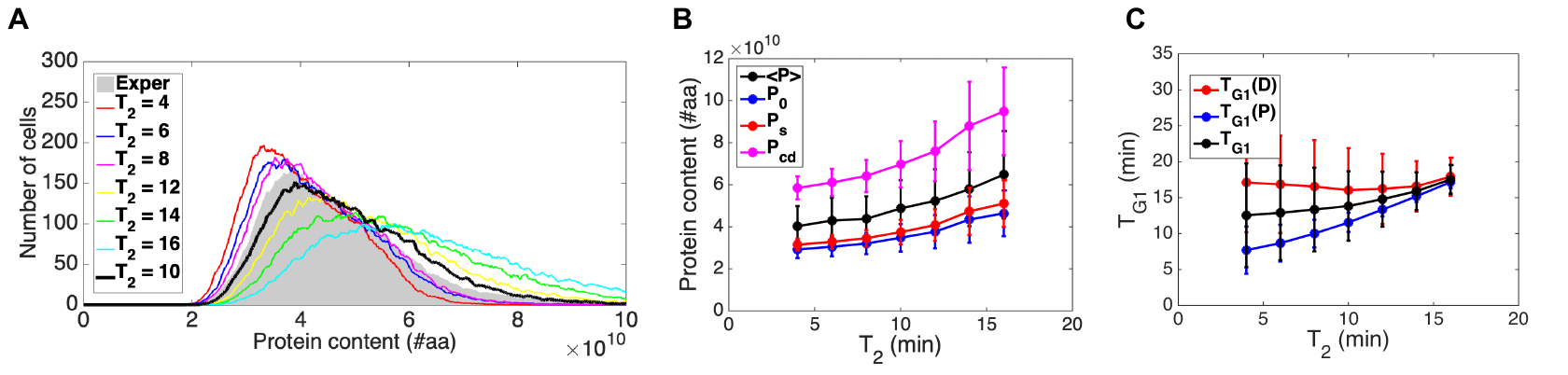

**Supplementary Figure S2 – Sensitivity analysis with respect to *T_2_* (glucose 2%)**. **A:** simulated protein distributions drawn for different values of *T_2_* and compared to the experimental one related to 2% glucose. **B:** protein features (average and initial cell protein content, critical cell size and size at division) extracted from the simulated populations of panel A, endowed with their standard deviations. **C:** G1 phase length for the whole population and for the subpopulations of daughters and parents, related to the simulated populations of panel A, endowed with their standard deviations.

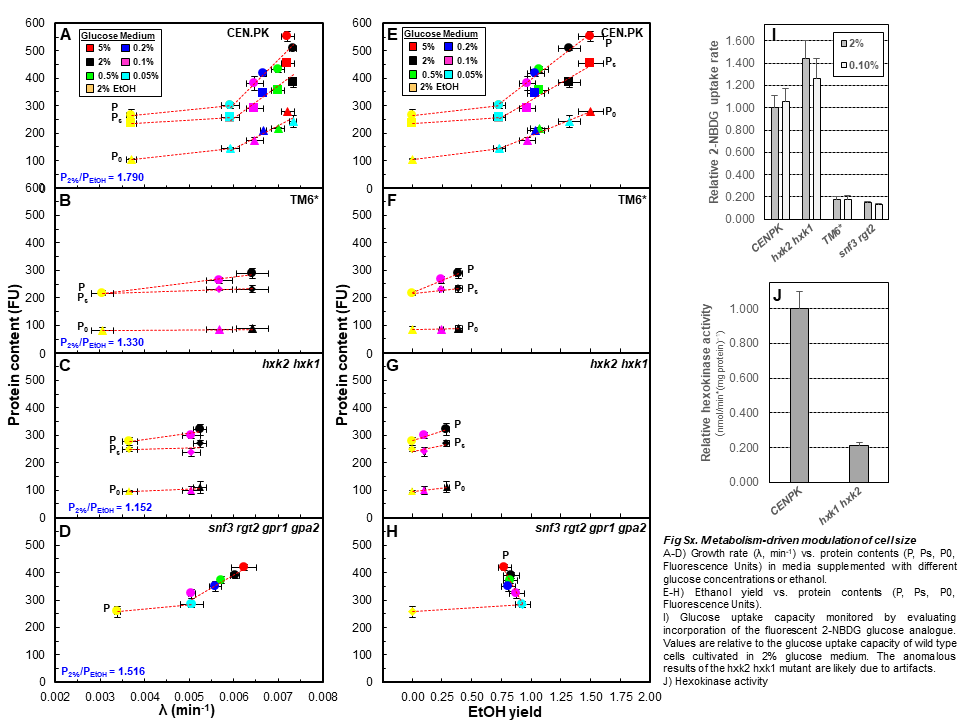
**Supplementary Figure S3**

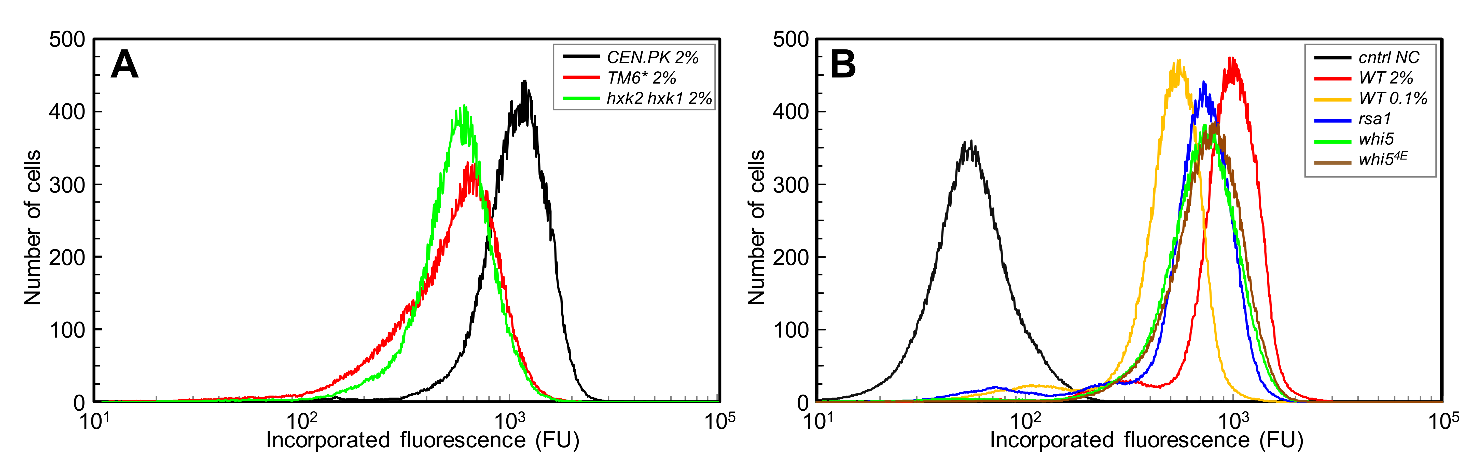

**Supplementary Figure S4 – Protein Synthesis rate**

Fluorescence incorporated in neosynthesised proteins was measured by cytofluorimetric analysis. Representative distribution profiles from various strains and growth conditions are shown.
